# New Statistical Metric for Robust Target Detection in Cryo-EM Using 2DTM

**DOI:** 10.1101/2024.10.01.616095

**Authors:** Kexin Zhang, Pilar Cossio, Aaditya Rangan, Bronwyn Lucas, Nikolaus Grigorieff

## Abstract

2D template matching (2DTM) can be used to detect molecules and their assemblies in cellular cryo-EM images with high positional and orientational accuracy. While 2DTM successfully detects spherical targets such as large ribosomal subunits, challenges remain in detecting smaller and more aspherical targets in various environments. In this work, a novel 2DTM metric, referred to as the 2DTM p-value, is developed to extend the 2DTM framework to more complex applications. The 2DTM p-value combines information from two previously used 2DTM metrics, namely the 2DTM signal-to-noise ratio (SNR) and z-score, which are derived from the cross-correlation coefficient between the target and the template. The 2DTM p-value demonstrates robust detection accuracies under various imaging and sample conditions and outperforms the 2DTM SNR and z-score alone. Specifically, the 2DTM p-value improves the detection of aspherical targets such as a modified artificial tubulin patch particle (500 kDa) and a much smaller clathrin monomer (193 kDa) in simulated data. It also accurately recovers mature 60S ribosomes in yeast lamellae samples, even under conditions of increased Gaussian noise. The new metric will enable the detection of a wider variety of targets in both purified and cellular samples through 2DTM.

## 1. Introduction

Accurately placing macromolecular assemblies in the cellular context is important in understanding their mechanistic role inside the cell. Previously, we developed a 2D template matching (2DTM) approach (Rickgauer *et al*., 2017; Lucas *et al*., 2021) in *cis*TEM (Grant *et al*., 2018) to detect targets in cellular cryo-EM images with high positional and orientational accuracy. 2DTM not only detects targets such as ribosomes in cryo-EM images but also provides data that enable the *in situ* classification and high-resolution reconstruction of these targets (Lucas *et al*., 2022; Lucas *et al*., 2023; Elferich *et al*., 2022). Building on these successes, this work aims to improve the 2DTM framework to detect more challenging targets in various environments.

A 2DTM search yields a signal-to-noise ratio (SNR) for every location in the cryo-EM image that depends on the cross-correlation between the template and the image (Rickgauer *et al*., 2017). A target is detected when the SNR value exceeds a statistically defined threshold that limits the average false positives to one per image, based on the assumption that the cryo-EM image is dominated by noise and cellular background and that the cross-correlation values observed across the image after whitening the noise/background follow a Gaussian distribution. The 2DTM SNR can be further normalized by subtracting the mean and dividing by the standard deviation of cross-correlations calculated across all sampled orientations at each location in the image (Rickgauer *et al*., 2017). This step is often referred to as “z-score” normalization (Spiegel & Stephens, 1999). Using the z-score instead of the SNR improves the detection of rotavirus double-layered particles (DLPs) (Rickgauer *et al*., 2017) and ribosomes in a crowded cellular environment (Lucas *et al*., 2022). In the following, we will refer to the outputs of 2DTM as 2DTM SNR and 2DTM z-score, respectively.

Previous applications of 2DTM have shown that the 2DTM SNR and z-score function differently depending on the characteristics of the sample and target. For example, when low-resolution features were suppressed by using a near-focus image setting (70 nm), the 2DTM SNR map showed a flat background with sharp peaks indicating the locations of apoferritins, even in a dense protein (BSA) background (Rickgauer *et al*., 2017). On the other hand, low-resolution features from the target itself when strongly defocused (*>* 2000 nm), or from the background structural noise, can result in broader peaks or an uneven background in the SNR map, complicating target detection (Rickgauer *et al*., 2017; Lucas *et al*., 2022). The misleading low-resolution background can be suppressed by calculating the 2DTM z-score (Rickgauer *et al*., 2017), which removes spurious correlations between the template and the structural noise in the image, thereby flattening the background and improving detectability of targets in cellular environments (Rickgauer *et al*., 2020; Lucas *et al*., 2022). In Fig. 1A, a segment of a previously published micrograph of a yeast lamella near the nucleus is presented (Lucas *et al*., 2022). This image section contains various cellular compartments located from left to right, including the vacuole, cytoplasm, and nucleus. Using the mature 60S as a search template, 2DTM outputs a 2DTM SNR map and a 2DTM z-score map (Fig. 1B and C). The bright spots in the 2DTM SNR map are locations with high correlation values, indicating 60S ribosomes. However, the peaks are surrounded by halos of increased SNR values extending to other low-resolution features in the image, such as membranes. The z-score map removes these halos and spurious matches of high-contrast features, thereby reducing the number of false detections (membranes or partial overlap with ribosomes) while preserving locations with high-resolution matches from the ribosomes.

**Fig. 1.**
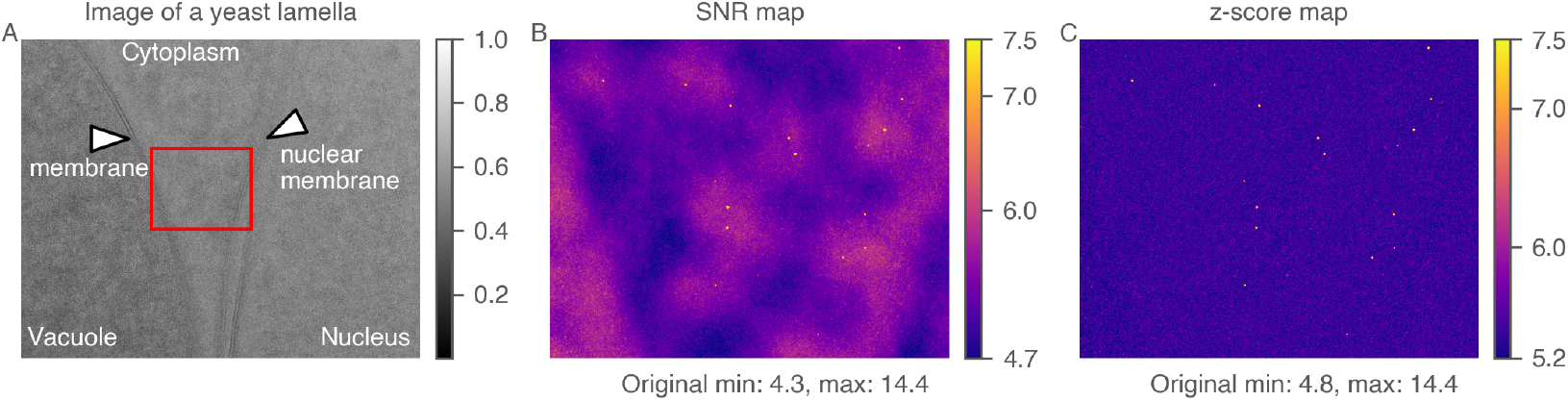
Comparison between the 2DTM SNR and the 2DTM z-score. (A) Micrograph section of a FIB-milled yeast lamella showing different compartments of the cell. (B) The 2DTM SNR map corresponding to the rectangle in (A), searched with a mature 60S template. (C) The 2DTM z-score map corresponding to the rectangle in (A). The pixel values were cropped to a narrower range (labeled on the color bars) for better visualization. The original data range is labeled below.

Despite its success, the current 2DTM workflow faces several challenges. Originally, the goal of 2DTM was the detection and localization of unlabeled molecules in the cellular context (Rickgauer *et al*., 2017), but more recent works have expanded its scope to other applications (Lucas *et al*., 2023; Lucas & Grigorieff, 2023). The question of detectability depends, therefore, on the target to be searched and the imaging conditions. First, in images of a purified sample, the likelihood that a low-resolution feature in the image is a valid target is high, making low-resolution contrast a reliable indicator of a true positive. However, this useful information is down-weighted in the z-score. Second, detecting smaller targets remains difficult since the crosscorrelation value depends on the size of the target. Although the detection limit of 2DTM was estimated as 150 kDa for purified samples (no molecular background) and 300 kDa for cellular samples (Rickgauer *et al*., 2017), no studies have systematically explored the detection of targets smaller than 200 kDa without incorporating prior information about their locations. Third, the shape of the targets plays a critical role in their detectability. Previous research has primarily focused on spherical targets, but as we demonstrate below, non-spherical shapes present additional challenges that have not been fully addressed. Finally, the detection threshold is based on the 2DTM SNR and images lacking strong low-resolution contrast. It remains unclear whether this threshold applies to other types of images or the 2DTM z-score. These factors highlight the need for further refinement of the 2DTM workflow to improve its applicability to a broader range of targets.

In this work, we investigate the performance of 2DTM applied to smaller and aspherical targets. We develop a novel metric, the 2DTM p-value, which combines information from the 2DTM SNR and z-score. We show that the 2DTM p-value has a more robust performance under varying imaging and sample conditions compared to using the 2DTM SNR or z-score alone. In particular, we demonstrate that the 2DTM p-value improves the detection of clathrin, a previously unexplored target due to its small size and aspherical shape, in simulated images under varying imaging conditions. We also show that the 2DTM p-value accurately recovers mature 60S ribosomes in yeast lamellae samples, even in increased levels of Gaussian noise.

## 2. Theory

Our current implementation of 2DTM outputs two scores, the 2DTM SNR and z-score. The novel 2DTM p-value integrates these two *features* into a new “metafeature”. The 2DTM p-value is designed to improve the detection of smaller and distinctly aspherical targets by utilizing correlations between the template and target across the entire resolution spectrum.

### 2.1. Previously developed metrics: 2DTM SNR and z-score

During a 2DTM search, we generate 2D projections from a 3D density map (*V*) of the molecule of interest (3D template) across over a million orientations (*τ*) within SO(3) space and multiply them with the contrast transfer function (CTF). A 2D projection is denoted as

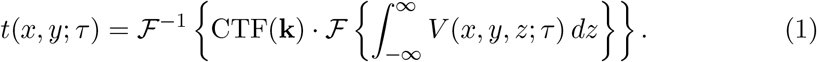

CTF parameters, such as defocus, can be estimated using the software package `CTFfind` (Rohou & Grigorieff, 2015; Elferich *et al*., 2024) and subsequently included in the search. We whiten the image to be searched, apply the same whitening filter to each 2D projection, then pad the 2D projection to the same size as the image. The whitened image *Y* and padded 2D projection *t*^*′*^(*i, j*; *τ*) are then normalized to zero mean and unity variance by

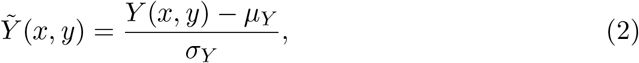

and

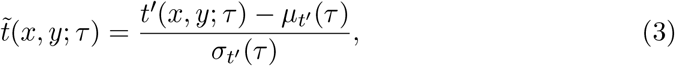

where *µ*_*Y*_, *σ*_*Y*_ are the mean and standard deviation of *Y*, and 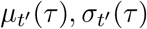 are those of *t*^*′*^(*i, j*; *τ*). We then calculate the cross-correlation for each 2D projection-image pair

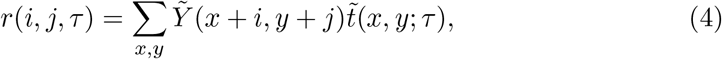

evaluated at all *i, j* locations in the image (Sigworth, 2004; Rickgauer *et al*., 2017). For each *i, j* location, we record the maximum cross-correlation

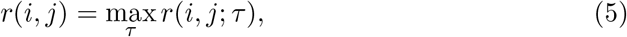

along with the best-aligned orientation

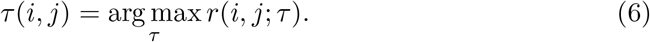

Additionally, the mean cross-correlation 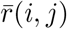, and the standard deviation of correlations *σ*(*i, j*) at each *i, j* location are calculated over sampled orientations as

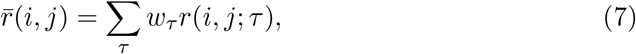

and

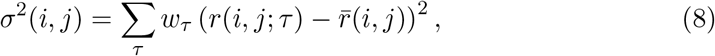

where *w*_*τ*_ is the quadrature weight to approximate integration in SO(3) space where ∑ _*τ*_ *w*_*τ*_ = 1.

The 2DTM *SNR* at location *i, j* in the image is then defined as the ratio of *r*(*i, j*) to the standard deviation of the cross-correlation values when only noise is present,

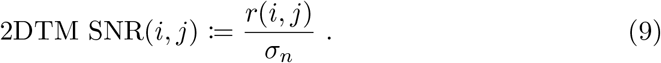

Assuming that the signal from detectable targets generates only a small amount of the variance of the entire image, we can estimate *σ*_*n*_ using the standard deviation of correlation values from the entire image

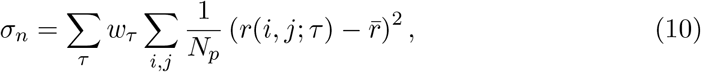

where *N*_*p*_ is the number of pixels in the image and 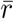 is the average of all the correlation values calculated in the search

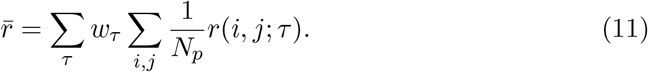

Given the normalizations of image and projection given in Eq. 2 and 3, previous work (Grigorieff, 2000) showed that

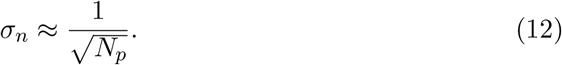

The 2DTM *z-score* at each *i, j* location is calculated by subtracting the average correlation from the maximum correlation and dividing by the standard deviation of the correlation values over the entire orientational space

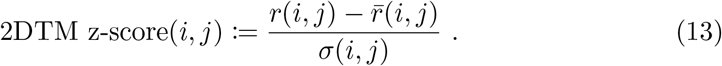

The 2DTM *targets* are generated by identifying local maxima in the 2DTM z-score map using a user-defined exclusion radius (typically 10 pixels). The targets are then subjected to a z-score threshold defined as

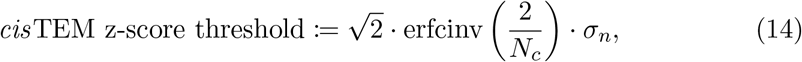

where erfcinv is the inverse complementary error function. In this equation, *N*_*c*_ is the number of correlation values calculated during the search. While factors such as symmetry in the template (as in the case of apoferritin) reduce the number of independent searches, leading to a z-score threshold that may be too high, we will show later that our new metric is unaffected by symmetry.

### 2.2. Theoretical analysis of the 2DTM z-score

In the following, we demonstrate that (a) the 2DTM z-score effectively removes the correlations originating from the rotationally invariant components of the template, and (b) the 2DTM z-score is related to the Fisher information which quantifies how tightly peaked the likelihood is at *r*(*i, j*) regarding orientation *τ*.

To prove the first point, we can write *r*(*i, j, τ*) as the sum of correlations from rotationally invariant component *r*_const_(*i, j*) and variant component *r*_vary_(*i, j, τ*). We can minimize the norm of the rotationally variant component *r*_vary_(*i, j, τ*) by setting the *τ* -average of *r*_vary_(*i, j, τ*) to zero

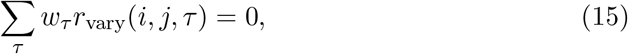

so that 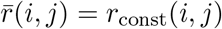. This implies that by constructing the z-score, the rotationally invariant components will be removed in Eq. 13. The rotationally invariant correlation components originate primarily from the low-resolution signal that is due to background structural noise that shares a similar size as the template. Previous studies have shown that Zernike polynomials can be used to decompose cryo-EM maps and analyze the continuous heterogeneity of biological macromolecules (Herreros *et al*., 2021; Herreros *et al*., 2023). We show in Appendix B that calculating the Zernike moments and Zernike invariants of a 3D template allows us to quantify the relative weight of the rotationally invariant and variant components, thereby measuring the asphericity of a template.

We next explore the relationship between the z-score and Fisher information regarding perturbations in *τ*. In previous works, Fisher information was applied to study how well a 3D potential map can be estimated from noisy, randomly rotated 2D projections under different noise levels (Fan *et al*., 2023; Fan *et al*., 2024). Here, we calculate the Fisher information regarding perturbations in *τ* instead. We first note that the l2-norm of the difference between the shifted image *Y* (*x, y*; *i, j*) and padded projection *t*^*′*^(*x, y*; *τ*) can be written as

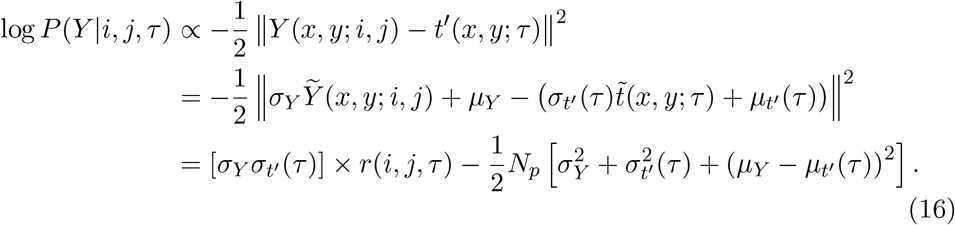

Assuming that each point-spread function has the same integral and the particle is not too aspherical, we have 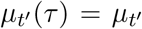 and 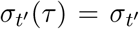. If we further assume that the image is generated by taking the “true” signal and adding independent identically-distributed pixel noise, we see that *r*(*i, j, τ*) is an affine transformation of the logarithm of the probability *P* (*Y* |*i, j, τ*) of observing the image *Y*, given a single particle at location *i, j*, and orientation *τ*,

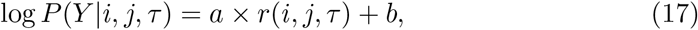

where 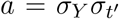and 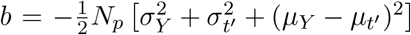. We see the Fisher information *I*(*τ*) is proportional to the second-*τ* -derivative of *r*(*i, j, τ*)

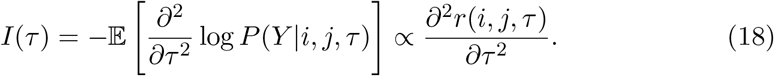

We consider the simple case of 1-dimensional *r*(*ψ*), where *ψ* is in domain Ω. *r*(*ψ*) can be roughly modeled as a Gaussian profile with a single peak (Appendix Fig. 12)

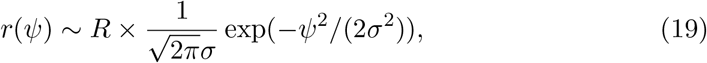

with zero mean, standard deviation *σ*, and a scaling factor *R*. We can calculate the following maximum (*r*_upb_), average (*r*_avg_), and variance 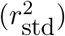 of *r*(*ψ*) regarding *ψ* as

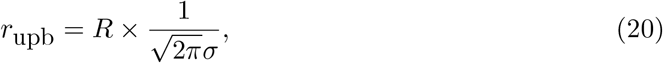

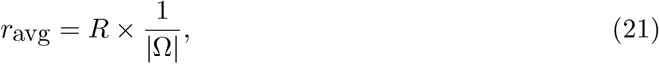

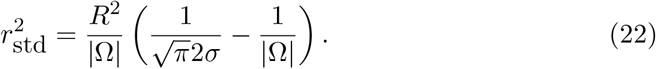

The 2DTM z-score, *z*, based on its definition, is

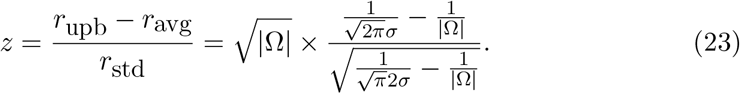

If we assume that the ‘size’ of the space |Ω| is relatively large compared to *σ*, then

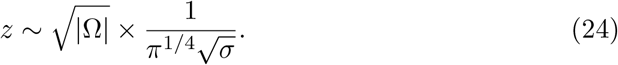

Meanwhile, the Fisher information of likelihood at *r*(*i, j*) with respect to perturbations in *ψ*, based on its definition in Eq. 18, is

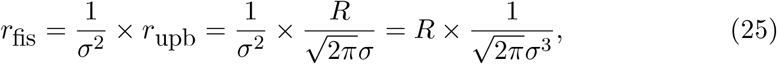

from which we could see that the z-score is related to the Fisher information by

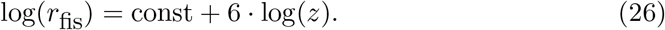

In Appendix A, we extend our discussion to *τ* ∈ SO(3) and demonstrate that, while the 2DTM SNR and z-score are related, they are not entirely redundant and can effectively complement each other. Specifically, peaks in the SNR map roughly correspond to local maxima in the log-likelihood of observing the data given that a true particle is being imaged (Eq. 17), whereas peaks in the z-score map correspond to locations and orientations with high Fisher information (Eq. 26). Given these differences, we aim to develop a method that integrates the 2DTM SNR and 2DTM z-score.

### 2.3. Quantile normalization

We now propose a general strategy for designing a “metafeature” that integrates the 2DTM SNR and z-score, estimating the probability of target detection without relying on a fixed z-score threshold. To combine features with varying scales, we first identify the local maxima in the 2DTM z-score map, extracting their corresponding 2DTM SNR and z-score values. Next, we independently apply a probit-function to both the z-scores and the SNRs (Amaratunga & Cabrera, 2001). The probit-function transforms the marginal distributions of both features into the standard Gaussian distribution, *N* (0, 1), with zero mean and unit variance. This method can be easily extended to more than two features and applied to datasets with even greater scale differences. The resulting quantile-normalized data is referred to as

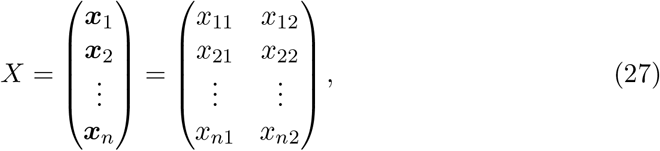

where ***x***_*i*_ ∈ ℝ^2^ is a 2D vector encoding the quantile-normalized features for a particular data point. For example, for the *i*-th data point ***x***_*i*_, *x*_*i*1_ represents the transformed 2DTM z-score and *x*_*i*2_ represents the transformed 2DTM SNR.

### 2.4. Fit with anisotropic Gaussian

To derive our new statistic, we fit an anisotropic Gaussian to the transformed data matrix *X*. This fit involves the empirical covariance matrix *C*^−1^, or equivalently the precision matrix *C*. We perform the eigenvalue decomposition of *C*^−1^ as

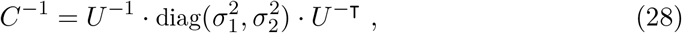

where diag 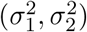 is the diagonal matrix formed from eigenvalues 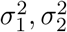, and the unitary matrix *U*^−1^ = [***u, u***_*⊥*_]. This results in an anisotropic Gaussian distribution fit to *X* with elliptical contours with major-axis ***u*** of length *σ*_1_ and minor-axis ***u***_*⊥*_ of length *σ*_2_. Our assumption below is that the joint distribution of *X* is well approximated by

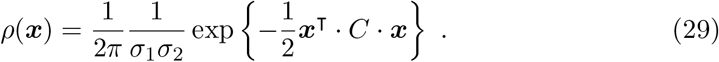

### 2.5. Calculation of the 2DTM p-value

Given a particle data point ***x***_*k*_, we can define *p*_*k*_, the probability of finding a sample from *ρ* in the first quadrant with a lower probability density than ***x***_*k*_, as

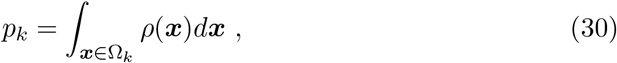

where the ‘first-quadrant” domain Ω_*k*_ (Fig. 2) is defined as

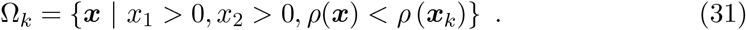

Describing the direction of ***u*** using angle *ω* such that ***u*** = [cos(*ω*), sin(*ω*)]^?^, *p*_*k*_ can be calculated by transforming the anisotropic Gaussian into a standard Gaussian and then integrating this standard Gaussian within the wedge corresponding to the (now transformed) first quadrant. The relevant angle associated with this wedge is

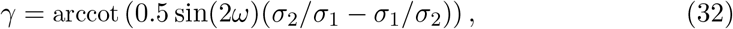

and *p*_*k*_ can then be written as

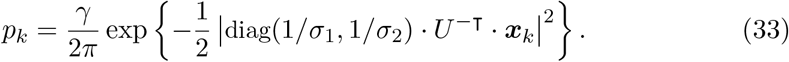

**Fig. 2.**
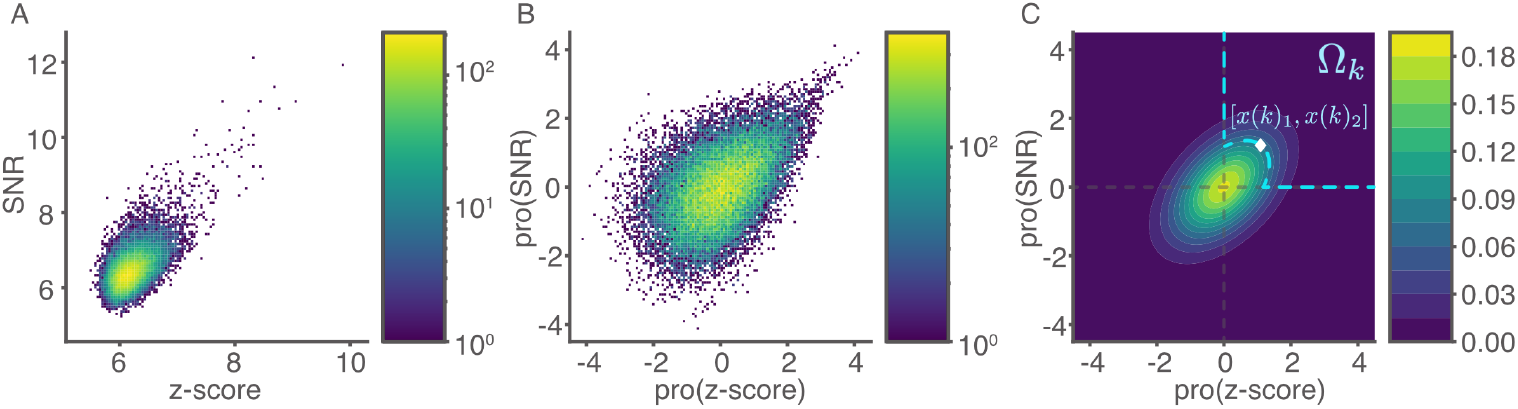
Computing the 2DTM p-value from the 2DTM SNR and z-score. (A) 2D histogram of the 2DTM SNRs versus z-scores for one of the clathrin montages (shown in Fig. 7A). (B) 2D histogram of the quantile transformed features in (A). Both (A) and (B) are plotted on a log scale. (C) Schematic plot for calculating the 2DTM p-value. A data point *x*(*k*) = [*x*(*k*)_1_, *x*(*k*)_2_], denoted by the diamond sign, represents the quantile-normalized data vector. The p-value is defined as the probability of finding a sample from the estimated anisotropic Gaussian that is rarer than *x*(*k*). Ω_*k*_ is the domain of samples to be considered that satisfies: (a) both transformed features should be larger than 0, and (b) the sample point should be rarer than *x*(*k*).

## 3. Results

### 3.1. Detection of simulated targets of distinct shapes in ice

To understand how molecular shape affects target detection by 2DTM, we simulated images of two molecules with similar molecular weights but distinct shapes and performed 2DTM searches on these images. The first target, apoferritin (Fig. 3A), is a spherical-shaped protein complex with octahedral symmetry that has frequently been used as a model system for benchmarking cryo-EM methods. In our study, we used the recently determined 1.27 Å resolution structure (PDBID: 7RRP), with a molecular weight of 498 kDa (Zhang *et al*., 2020). The second target is an artificial tubulin patch (different views shown in Fig. 3B and C) derived from the single-particle model of deacetylated microtubules (PDBID: 6O2S) (Eshun-Wilson *et al*., 2019). This rod-shaped particle consists of six *α*-tubulin subunits, four *β*-tubulin subunits, and one additional modified *β*-tubulin subunit, with a total molecular weight of 500 kDa.

**Fig. 3.**
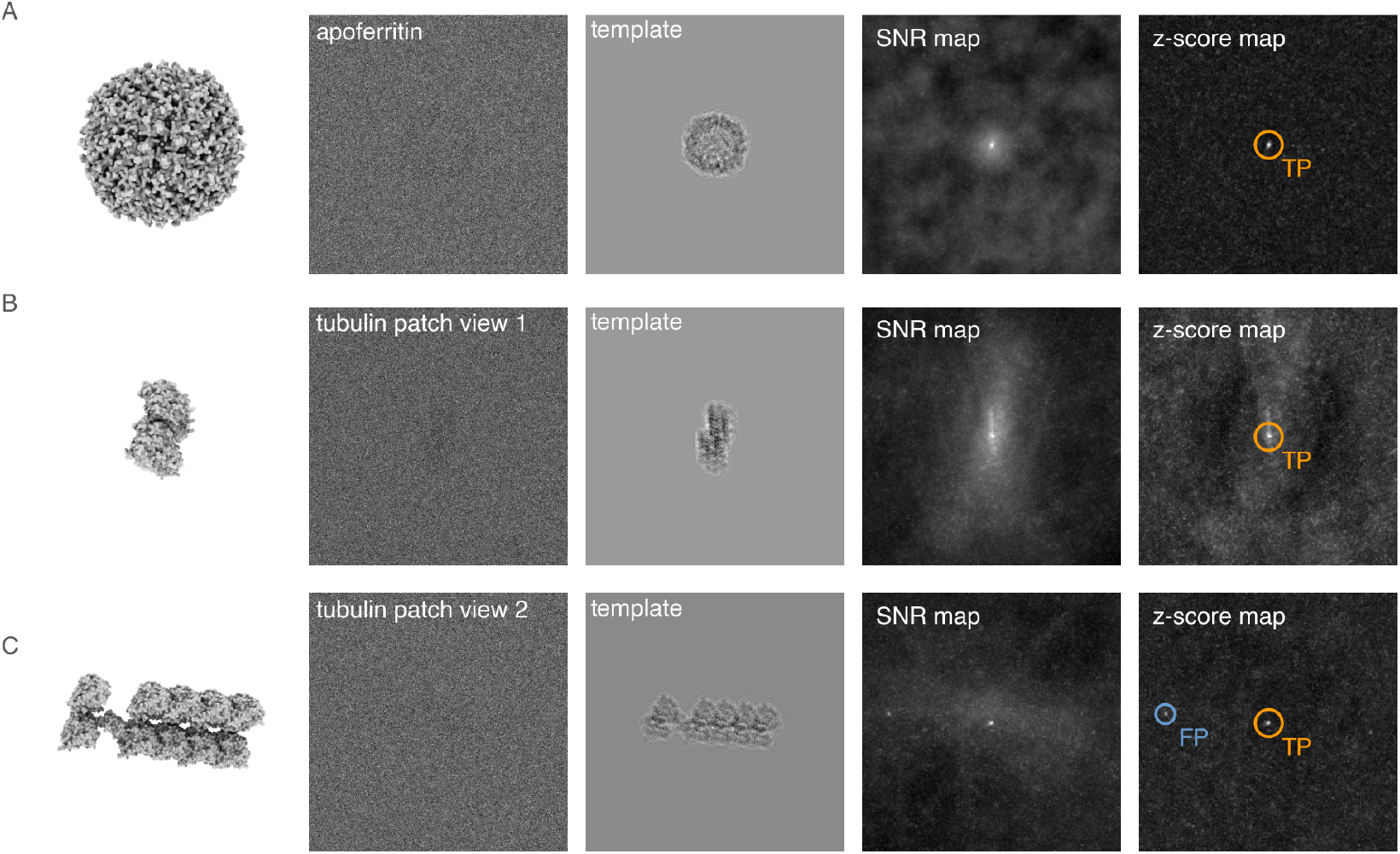
Local 2DTM SNR and z-score maps of an apoferritin particle (A) and two tubulin patch particles (B-C). Each row shows the 3D template generated from *cis*TEM program `simulate` (with similar views as the particles), the simulated particle at an underfocus of 500 nm, the 2DTM matched template, the SNR map, and z-score map centered on the particle. Targets that survived the *cis*TEM z-score threshold are circled. True positives (orange) are located near the center of the particles. Templates in rows (B) and (C) represent the tubulin patch viewed edge-on and from the side.

The simulated particle images at different orientations were generated using the program `simulator` in *cis*TEM (Himes & Grigorieff, 2021), with an underfocus of 500 nm and a uniform B-factor of 30 Å^2^ (Fig. 3 second column). The simulations were performed with a pixel size of 1.0 Å and a total exposure of 30 e^−^/Å^2^ in 100 nm ice. Segments of the 2DTM SNR and z-score maps of the three simulated particle images are shown in the third and fourth columns. Targets exceeding the *cis*TEM z-score threshold were labeled as either true positives (orange) or false positives (blue). Similar to the 60S ribosome (Fig. 1), the z-score map of the apoferritin particle features a sharp peak with a clean background (Fig. 3A). In contrast, the z-score map of the tubulin patch particle is either noisier (Fig. 3B) or contains a false positive (Fig. 3C).

We then simulated cryo-EM images of 100 apoferritin particles and 100 tubulin patch particles, each with defocus values of 70 and 2000 nm (Rickgauer *et al*., 2017), in random orientations. We arranged the 100 particle images into a pseudo cryo-EM image in a 10-by-10 montage. A segment of the montage containing four tubulin patch particles (arranged 2-by-2) at 2000 nm defocus is shown in Fig. 4A. We searched the montages by 2DTM using an angular search grid with an in-plane step of 1.5^°^ and an out-of-plane step of 2.5^°^. The defocus was searched in a range of +/-100 Å and a step size of 20 Å (Lucas *et al*., 2021). 2DTM *targets* were identified as local maxima within a 10-pixel radius in the z-score map and labeled according to their angular and translational errors relative to the expected values from the simulation, taking into account the octahedral symmetry of apoferritin. The angular error was calculated based on the average l2 distances between points in the two (unit vector) templates after angular transformation. The translational error (*d*_*xy*_) was defined as the distance between the target and the grid center of the closest simulated particle. The distribution of errors for one of the tubulin patch montages at 2000 nm is shown in Appendix Fig. 13. We used a cutoff of 7 pixels for *d*_*xy*_ and 0.4 for Euler error for labeling the targets. True targets (orange) were accurately located near the centers of the particle grid cells, whereas false targets (blue), which resulted from partial overlaps with tubulin patches or matches with background noise, were not necessarily near the centers (Fig. 4B).

**Fig. 4.**
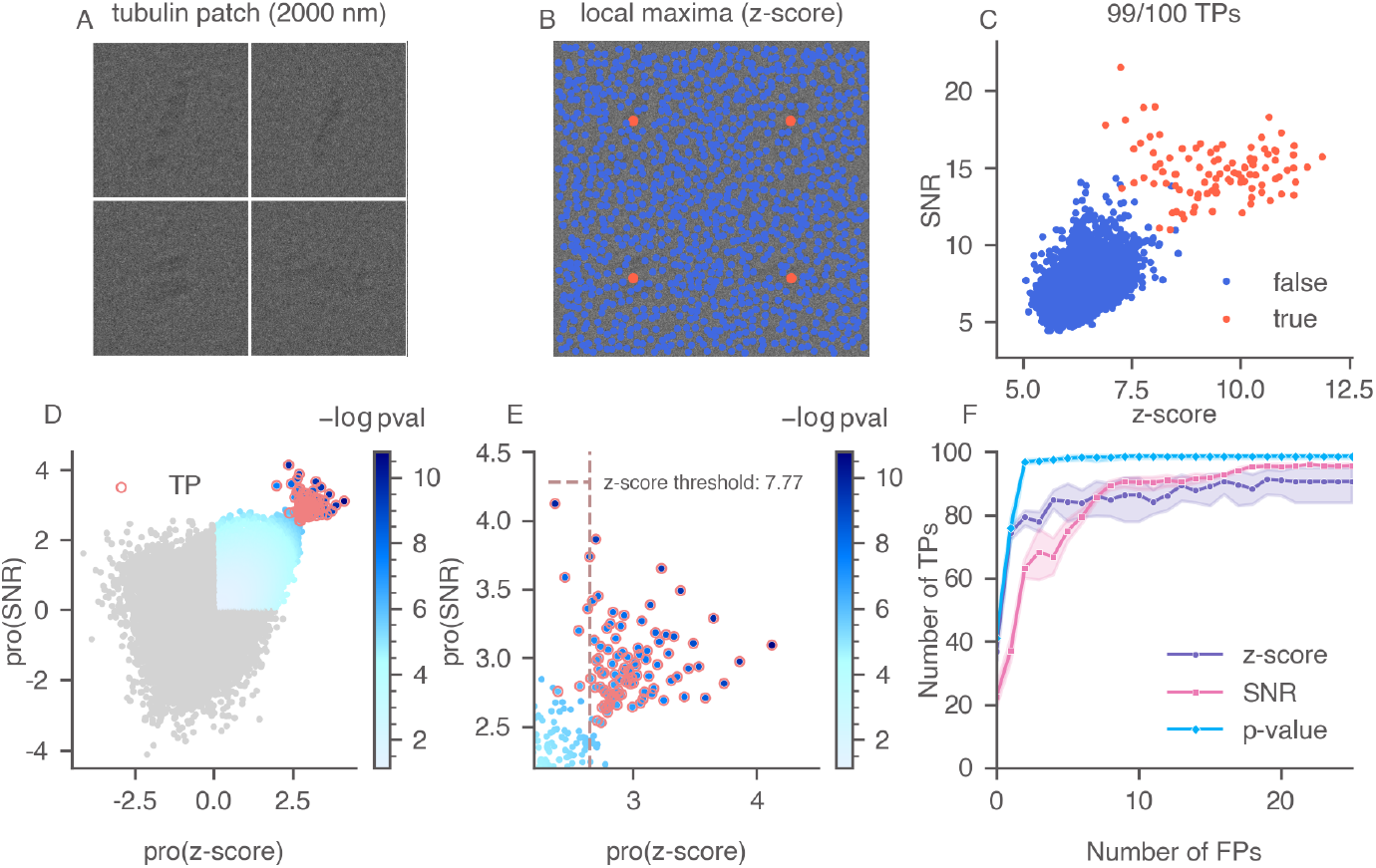
Calculation of the 2DTM p-value and evaluation of three 2DTM metrics for a simulated tubulin patch montage. (A) An example of a tubulin patch montage segment with four particles arranged in a 2-by-2 grid. (B) Local maxima identified in the 2DTM z-score map of the montage segment in (A) using a 10-pixel radius. (C) A scatter plot of 2DTM SNR versus z-score, with true targets labeled in orange based on their expected location and orientation. 99 of the 100 targets in the example montage were recovered as local maxima in the z-score map. (D) The quantile-normalized data, color-coded by their 2DTM p-values (− log(p-value)). Data points not in the first quadrant are labeled in gray, as they are excluded from the p-value calculation. True targets are circled in orange. (E) A zoomed-in version of (D) showing the transformed z-score threshold. (F) ROC curves of the three 2DTM metrics, with shaded areas indicating the confidence intervals calculated from 300 simulated particles. All the figures in this work were styled using the Python library `niceplots` (Gray *et al*., 2024).

Comparing the 2DTM SNRs to the z-scores for targets identified in the simulated montage (Fig. 4C), we found that using either feature individually led to a higher number of false positives or false negatives compared to using both features together. We applied quantile normalization to the 2DTM SNRs and z-scores of the targets and calculated the 2DTM p-values for data points located exclusively in the first quadrant after transformation (Fig. 4D), as a true target is expected to exhibit both high SNR and high z-score. The zoomed-in scatter plot (Fig. 4E) shows the transformed threshold corresponding to a *cis*TEM z-score threshold of 7.77. Several true targets fell below this threshold, indicating false negatives, while several false positives were observed near the threshold. We combined the results from 300 simulated particles and evaluated the accuracies by calculating their receiver operating characteristic (ROC) curves (Fig. 4F). To better understand the classification accuracy when a low false positive rate (FPR) is desired, we focused on a specific FPR range with fewer than 25 false positives. Our results show that the 2DTM p-value successfully recovered more true targets than the other two metrics.

Next, we compared the detection of apoferritin and tubulin patch at different defocus. In the 70 nm montages (Fig. 5A and 5G), particles were barely visible, whereas in the 2000 nm montages (Fig. 5D and 5J), there was strong low-resolution contrast from the particles. Analyzing the scatter plots of 2DTM SNRs versus z-score and the ROC plots, we made the following observations. First, at low defocus, the SNR values of the true targets showed strong correlations with the z-scores, in contrast to the correlations observed at higher defocus. Specifically, the Pearson correlations observed at 70 nm defocus were 0.92 for apoferritin (Fig. 5B) and 0.85 for tubulin patches (Fig. 5H). However, at 2000 nm defocus, these correlations dropped to 0.47 (Fig. 5E) and -0.19 (Fig. 5K), respectively. This is because, at low defocus, low-resolution contrast was suppressed, causing the 2DTM SNR to contain mostly high-resolution information, similar to the z-score. Second, for tubulin patches, the SNRs and z-scores of the true targets were less correlated compared to apoferritin, suggesting the two metrics may provide complementary information for aspherical targets. This finding highlights the need to design a “metafeature” that integrates both metrics, which our new metric, the 2DTM p-value, achieves. Finally, we found that for apoferritin, all three metrics showed comparable target detection accuracies at both defocus values, although the z-score exhibited slightly lower accuracies at low FPR ranges (Fig. 5C and 5F). In contrast, for aspherical particles, the p-value proved to be the optimal metric, with its performance improving as the low-resolution contrast from the particles increased, outperforming the z-score (Fig. 5I and 5L). Unlike the z-score threshold, the p-value is unaffected by template symmetry because it uses the statistics at the optimal orientation once it is found.

**Fig. 5.**
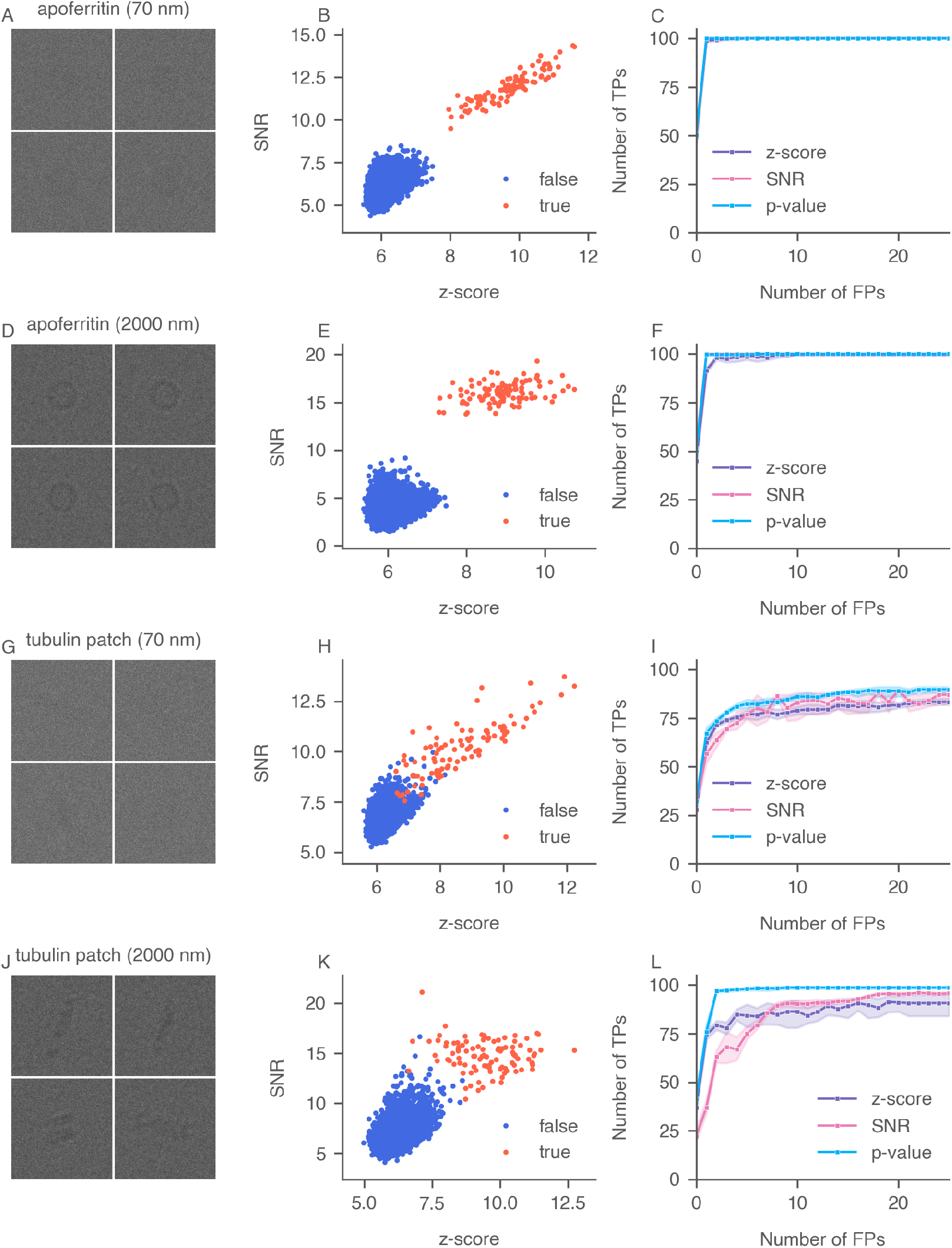
Comparison of 2DTM searches for apoferritin (A-F) and tubulin patch (G-L) at 70 nm and 2000 nm defocus. Each row shows a segment of the 10-by-10 montage, an example scatter plot of 2DTM SNR versus z-score, and the ROC curves of the three metrics calculated from 300 simulated particles.

### 3.2. Detection of simulated clathrin monomers in isolation

We next focused on the more difficult task of detecting both small and aspherical targets. Clathrin is a protein crucial for endocytosis, facilitating the cellular uptake of substrates from the extracellular environment (Kaksonen & Roux, 2018). It forms a three-dimensional lattice known as a clathrin coat, which transports vesicles with cargo to be endocytosed. The clathrin triskelion consists of three heavy chains that interact at their C-termini, with each heavy chain tightly bound to a nearby light chain (Fotin *et al*., 2004). The high-resolution structure of the invariant hub, determined using single-particle cryo-EM (PDBID: 6SCT), exhibits C3 symmetry (Morris *et al*., 2019). The template we used is the clathrin monomer, consisting of three heavy chains and two light chains, with a molecular weight of 193 kDa (orange part Fig. 6). Although the clathrin monomer slightly exceeds the previously reported detection limit of 2DTM (150 kDa for particles embedded in ice (Rickgauer *et al*., 2017)), it is the smallest target studied by 2DTM so far. Its relatively small molecular weight and highly aspherical shape provided an excellent test case for exploring the limits of 2DTM.

**Fig. 6.**
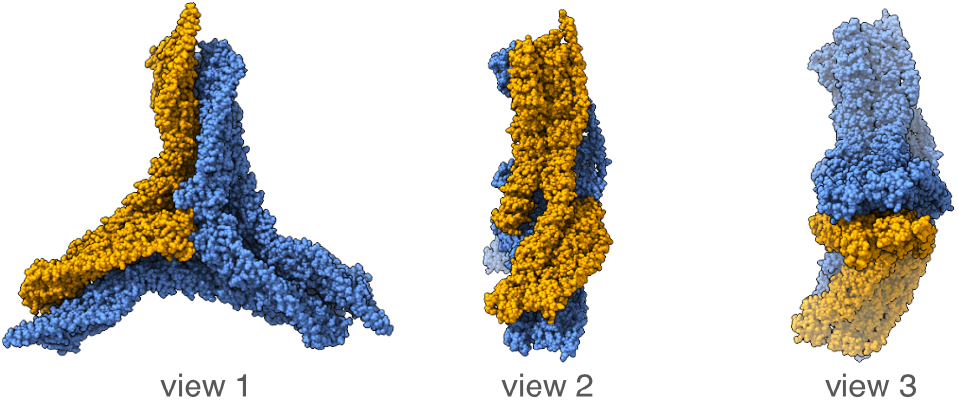
Structure of the clathrin monomer used as the template for 2DTM searches. Shown are different views of the complete clathrin invariant hub (blue), determined from single-particle cryo-EM, and the clathrin monomer (orange) used as the template in our experiments. The monomer consists of three heavy chains and two light chains, with a molecular weight of 193 kDa.

We simulated cryo-EM images of clathrin monomer particles in 100 nm ice in random orientations using the B-factor from the PDB entry, at a pixel size of 1.06 Å, arranged them into a pseudo cryo-EM image (a segment shown in Fig. 7A), and performed the 2DTM searches. Due to the smaller weight of the monomer, we increased the total dose to 45 e^−^/Å^2^ and used a defocus of 500 nm in simulation. A defocus search was performed on all images with a step size of 200 Å in a total range of 2400 Å (+*/*−1200 Å). 2DTM *targets* were identified as local maxima in the z-score map and labeled based on whether they were true (orange) or false (blue) (Fig. 7B). Out of the 100 simulated particles, 92 were recovered as local maxima. The overlap between the true and false positive populations (Fig. 7C) made it challenging to classify the targets by a binary threshold based solely on the z-score or the SNR.

**Fig. 7.**
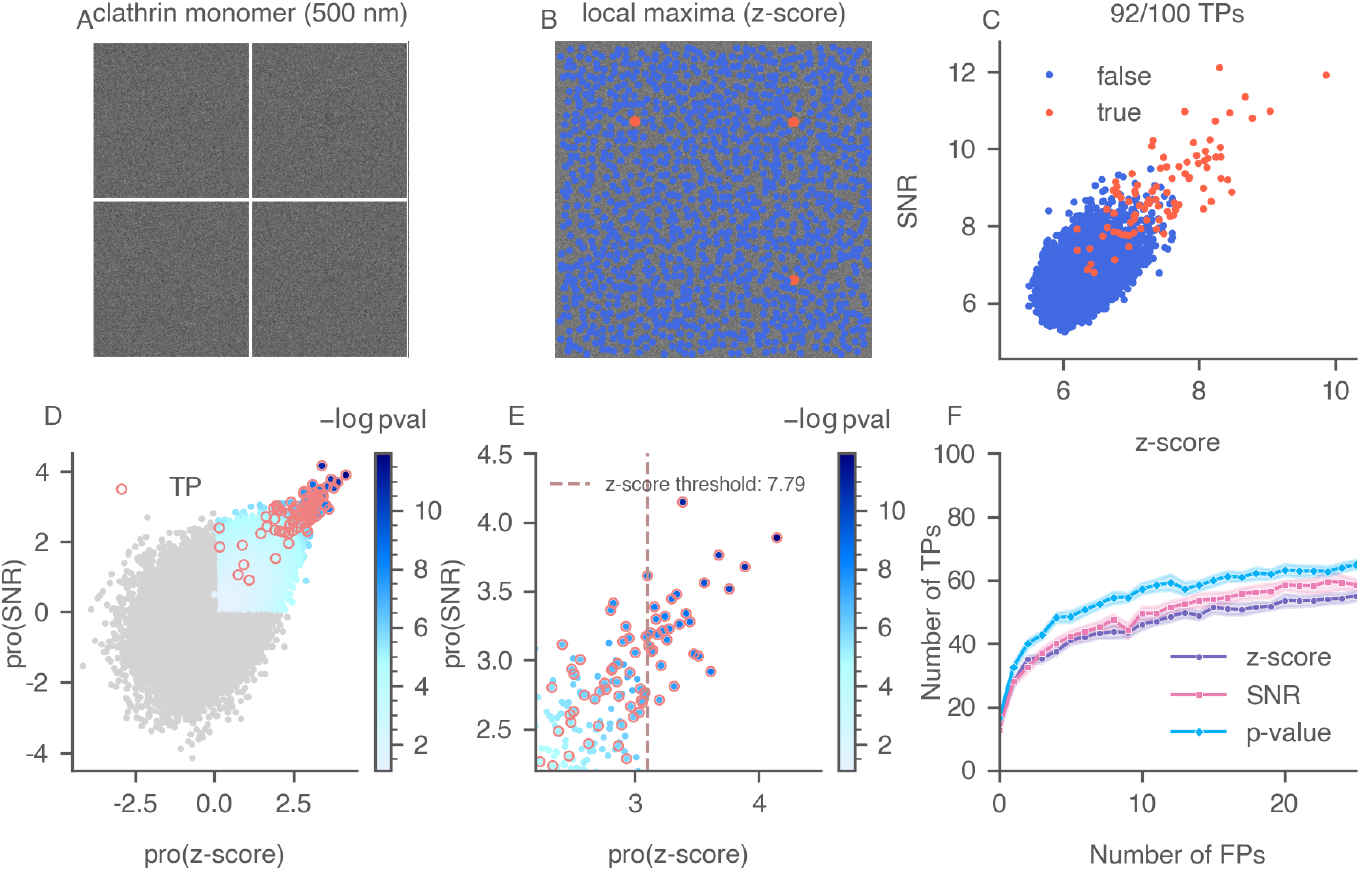
Calculation of the 2DTM p-value and evaluation of three 2DTM metrics for a simulated clathrin montage. (A) An example of a clathrin montage segment with four particles arranged in a 2-by-2 grid. (B) Local maxima identified in the 2DTM z-score map of the montage segment in (A) using a 10-pixel radius. (C) A scatter plot of 2DTM SNR versus z-score, with true targets labeled in orange based on their expected location and orientation. In total, 92 out of the 100 targets in the example montage were recovered as local maxima in the z-score map. (D) The quantile-normalized data, color-coded by their 2DTM p-values (− log(p-value)). Data points not in the first quadrant are labeled in gray, as they are excluded from the p-value calculation. True targets are circled in orange. (E) A zoomed-in version of (D) showing the transformed z-score threshold. (F) ROC curves of the three 2DTM metrics, with shaded areas indicating the confidence intervals calculated from 2000 simulated particles.

Compared to apoferritin and tubulin patch, detecting clathrin monomers using the *cis*TEM z-score threshold resulted in more false negatives (Fig. 7E), due to their smaller size. We repeated this analysis for 2000 simulated clathrin monomers and reported the performance of the three 2DTM metrics (Fig. 7F). Using the SNR instead of the z-score recovered more true positives at higher FPR levels, owing to its incorporation of low-resolution signal. In the FPR range with fewer than 25 false positives, the 2DTM p-value consistently outperformed the SNR and z-score, as it recovered more true positives for a given FPR, even under conditions where no false positives were allowed.

### 3.3. Detection of simulated clathrin monomers with increasing solvent background

Next, we examined how increasing solvent thickness, and consequently the solvent noise, affects the detection accuracy of the 2DTM p-value, particularly for smaller and more aspherical targets. The 2DTM SNR theoretically increases with the molecular weight of the template, limiting current 2DTM detection to around 150 kDa in ice and 300 kDa in 100 nm thick samples with protein background (Rickgauer *et al*., 2017; Rickgauer *et al*., 2020). To study targets in their native state, focused ion beam (FIB)-milling is used to cut sections (lamellae) of frozen-hydrated biological specimens. Typical lamella thicknesses range from 85 to 250 nm (Lam & Villa, 2021). However, increased sample thickness leads to the loss of electrons due to inelastic and multiple scattering, reducing image signal, particularly at higher resolution (Peet *et al*., 2019; Dickerson *et al*., 2022). Earlier studies have demonstrated the importance of correctly modeling the hydration layer in cryo-EM images (Shang & Sigworth, 2012; Himes & Grigorieff, 2021) and revealed an exponential decay in the 2DTM z-score with an increase in solvent thickness, particularly for detecting large ribosomal subunits (Rickgauer *et al*., 2020; Lucas & Grigorieff, 2023). However, the relationship between 2DTM detection accuracy and solvent thickness remains unexplored for targets less spherical than large ribosomal subunits. Therefore, we conducted simulations of clathrin monomers, systematically varying the solvent thickness within a range consistent with typical FIB-milled lamellae.

We examined three ice thickness values: 120 nm, 150 nm, and 200 nm. For each thickness level, we simulated 1000 clathrin monomers and created ten 10-by-10 montages (example montages in Fig. 8). The accuracy of all three 2DTM metrics declined with increasing solvent thickness due to the loss of high-resolution signal. Notably, the 2DTM p-value consistently outperformed the other metrics across diverse solvent conditions, although its performance converged with the SNR as the solvent thickness approached 200 nm.

**Fig. 8.**
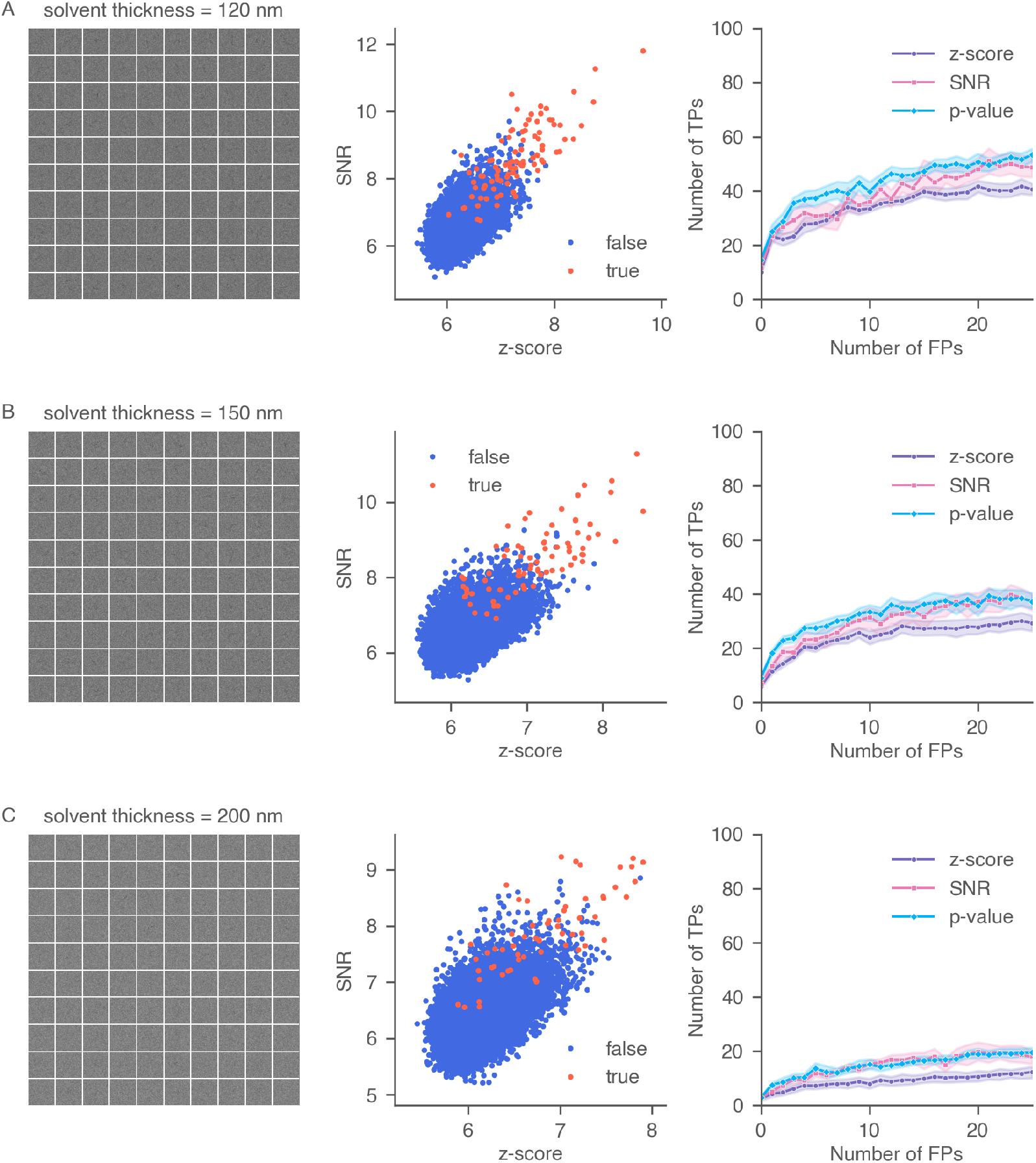
Performance of 2DTM metrics under different solvent thicknesses: (A) 120 nm, (B) 150 nm, and (C) 200 nm. For each thickness, the figures show (from left to right): a simulated montage containing 100 clathrin monomers, a scatter plot comparing 2DTM SNR and z-score, and ROC curves of the three metrics for 1000 simulated particles.

As previously discussed, the z-score depends heavily on the correlations between the image and the rotationally variant components of the template. The rotationally variant components tend to represent higher resolution than the invariant parts. In contrast, the SNR relies on both high- and low-resolution signal from the template. High-resolution signal decays faster than lower-resolution signal when ice is thick; therefore, the detection mainly depends on the low-resolution features.

### 3.4. Detection of clathrin monomers in simulated protein mixture images

Previous works have pointed out that the 2DTM SNR is more affected by the presence of proteins or structural features in the cell that share a similar size and shape as the target, making accurate detection more difficult (Rickgauer *et al*., 2017; Lucas *et al*., 2022). Since the 2DTM p-value is derived from the SNR, we explored in this section whether the 2DTM p-value can correctly detect clathrin monomers in images containing other proteins.

To simulate images with protein mixtures, we prepared montages, each containing 50 clathrin monomers and 50 proteasome particles (PDBID: 7LS6 (Schnell *et al*., 2021), 408.62 kDa) in random orientations. The clathrin monomer was used as the template in the 2DTM search. We simulated ten protein mixture montages (example in Fig. 9A) and compared the targets’ 2DTM SNRs against z-scores. The 2DTM p-value recovered more true positives compared to the 2DTM SNR and z-score, consistent with the results observed in images containing only clathrins. The z-score performed better than the SNR as it relies less on low-resolution signal that may come from incorrect particles. Target detection based solely on the 2DTM SNR led to false positives located near proteasome particles due to their stronger low-resolution contrast (Appendix Fig. 14A).

**Fig. 9.**
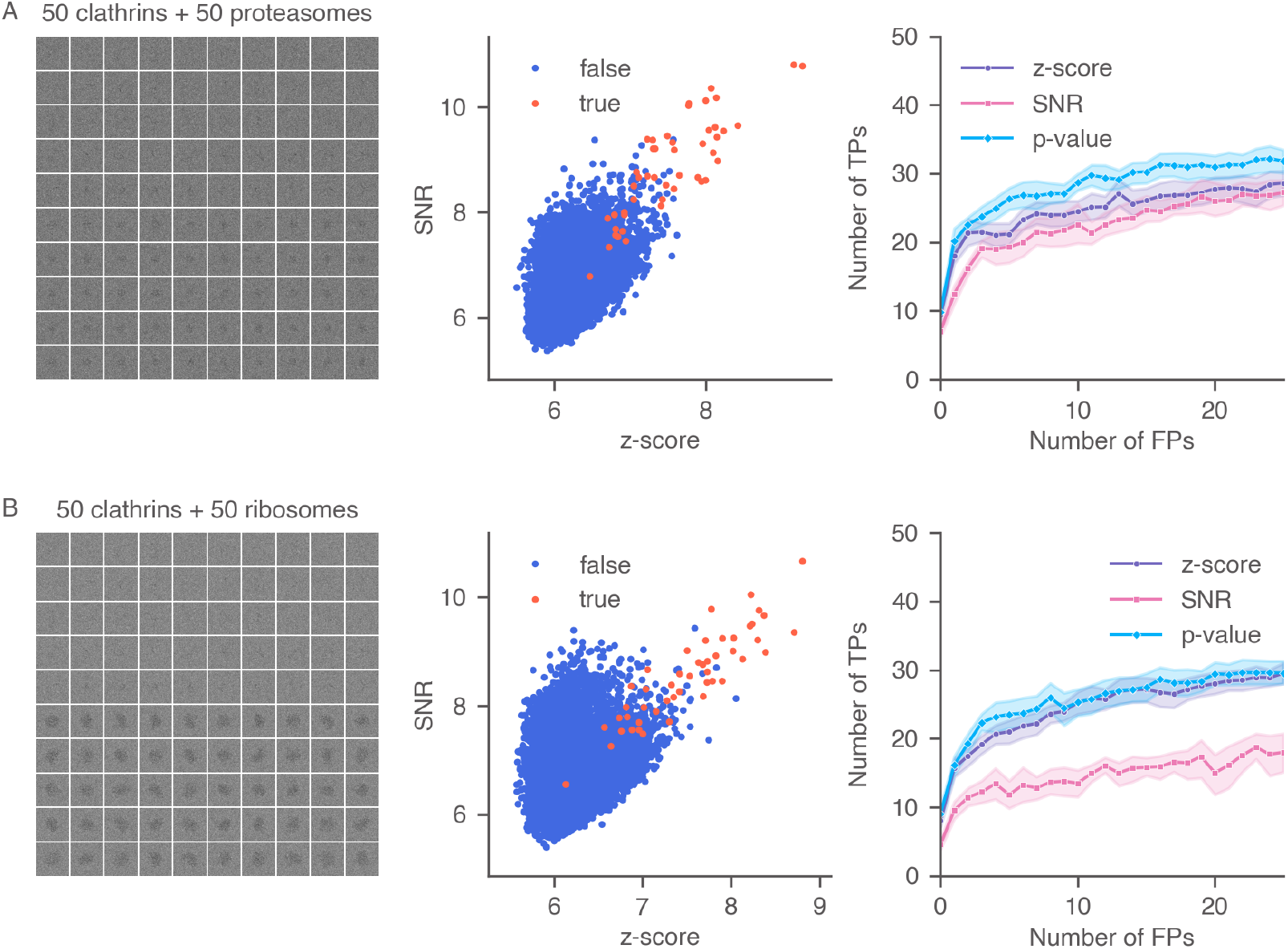
Performance of 2DTM metrics when searching a mixture particle montage using the clathrin monomer as the template. For each row, the figures show (from left to right): an example mixture montage containing 50 clathrin monomers and 50 proteasomes (A) or 50 mature 60S ribosomes (B), a scatter plot comparing 2DTM SNR and z-score, and ROC curves of the three metrics for ten 10-by-10 mixture montages (which correspond to 500 simulated clathrin particles mixed with 500 proteasome particles (or 60S particles)).

As a further test, we present an extreme scenario where we simulated ten protein mixture montage (example in Fig. 9B) containing clathrin monomer and mature 60S ribosomal subunits (PDBID: 6Q8Y (Tesina *et al*., 2019), 1.72 MDa) and searched using the clathrin monomer as the template. Due to the much stronger low-resolution signal of the 60S subunits, the 2DTM SNR was significantly higher at these locations, leading to incorrect detections. For the same FPR, the 2DTM SNR recovered significantly fewer true targets than the 2DTM z-score. Most locations with high SNR values were near the mature 60S particles (Appendix Fig. 14B). Interestingly, the 2DTM p-value still recovered as many or more clathrins compared to the z-score. This example confirms that even if the 2DTM SNR is unreliable in the presence of high background noise or other proteins, the performance of the p-value is similar to, if not better than, using the z-score alone. This highlights the potential of using the 2DTM p-value to study densely populated cellular images.

### 3.5. Detection of mature 60S in experimental images with added Gaussian noise

Next, we evaluated whether the 2DTM p-value maintained its superior performance in a setting where the z-score also performs well. Furthermore, we wanted to investigate the performance of the 2DTM p-value for target detection in experimental cryo-EM images of cellular lamellae. Given the lack of ground truth labels for experimental data, we introduced Gaussian noise with increasing variances to previously analyzed yeast lamellae (Lucas *et al*., 2022). We performed 2DTM searches using the mature 60S template (PDBID: 6Q8Y (Tesina *et al*., 2019)) and compared the detection results based on the 2DTM z-score, SNR, and p-value to those from images without additional noise. We systematically varied the ratio of added noise variance relative to the image variance, ranging from 0.1 to 2.5. Under each noise condition, we generated nine images featuring random Gaussian noise.

These images were selected because they contained a significant number of matches with high 2DTM z-scores, indicating high confidence in the identities of these locations. The small differences in the estimated defocus values between the noisy images and the corresponding Gaussian-noise-free images were within the 2DTM defocus search step and could, therefore, be ignored here. Comparing the images pre- and post-noise addition (Appendix Fig. 15 to 17), we observed that the high-resolution signal was gradually lost as the level of noise increased.

We next explain the manual labeling process using the pre-noise micrograph in Fig. 10A as an example. The thickness of this image was estimated to be 98 nm, and the average defocus was around 367 nm (Lucas *et al*., 2022). The relatively lower defocus suppressed low-resolution noise from the cellular background and improved the detection of the mature 60S using the 2DTM z-score. We calculated a threshold in the z-score histogram (Fig. 10B) that established a clear separation between *tail* scores (considered true positives) and *bulk* scores (considered true negatives), minimizing the overlap between the two. For locations in the image that do not contain mature 60S signal, their 2DTM z-score values should follow a *generalized extreme value* (GEV) distribution (Haan & Ferreira, 2006) as explained in Appendix C. The GEV distribution was superimposed on the histogram (dashed blue curve in Fig. 10B), fitted using z-scores smaller than the *cis*TEM z-score threshold (7.85). While the bulk z-scores were well modeled by the GEV distribution, the tail z-scores were not. The fitted distribution approached zero rapidly at around 8.0, while the tail of the histogram extended to 15.1 (inset of Fig. 10B), indicating strong correlations with the mature 60S template. Using the fitted GEV distribution as the null hypothesis, we calculated the 2DTM z-score corresponding to a given FPR. In this experiment, we set the FPR to 10^−6^, resulting in a z-score threshold of 8.212, consistent with a visual separation of the tail from the bulk. Targets were restricted from being detected within 100 Å to the edge of the image to avoid the detection of partial particles. 149 targets with z-scores exceeding 8.212 were labeled as true targets and plotted onto the micrograph based on their 2DTM-derived locations and orientations (Fig. 10A). Using the same strategy, we labeled two other images from yeast lamellae and found 176 and 336 true targets, respectively (Appendix Fig. 18 and 20).

**Fig. 10.**
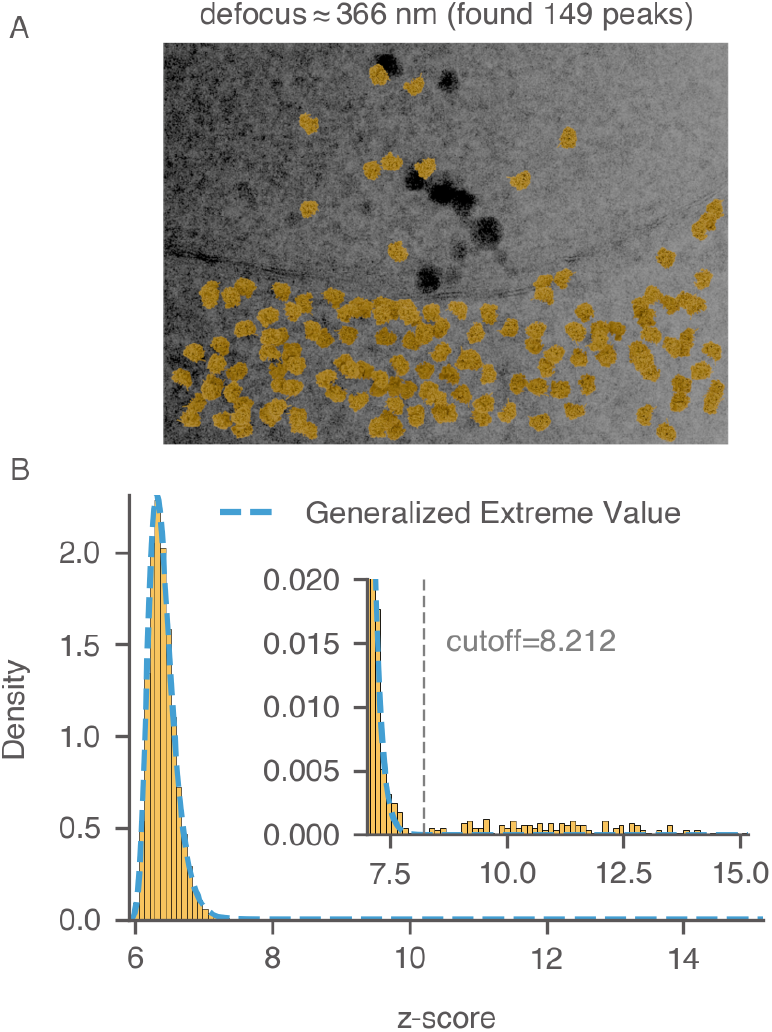
Labeling 60S targets in yeast lamella. (A) Image `150_Mar12_12.28.45_165_0.mrc` from previous work (Lucas *et al*., 2022). Control peaks are selected based on the threshold determined in (B). Particles are plotted with their 2DTM-derived alignment parameters. (B) Distribution of z-scores across all locations in the image using mature 60S as the template. The dashed blue curve represents the fitted generalized extreme value distribution. The threshold that best separates false matches (bulk) from true matches (tail) is labeled.

We calculated the detection accuracies of the 2DTM SNR, z-score, and p-value for the three lamellae upon adding varying levels of added Gaussian noise (Fig. 11, Appendix Fig. 19, and 21). In cases where minimal noise was introduced, particularly when the ratio of the added noise to the image noise was less than or equal to 0.5 ((Var(*n*)*/*Var(*I*) *≤* 0.5)), the performance of the 2DTM p-value generally aligned with that of the z-score and was better than that of the SNR. Since the control targets were labeled using the z-score, the z-score was expected to exhibit optimal accuracy under conditions of minimal Gaussian noise addition. For these three lamellae, when the noise ratio was 0.5, the 2DTM p-value started to outperform the z-score across most of the relevant FPR ranges.

**Fig. 11.**
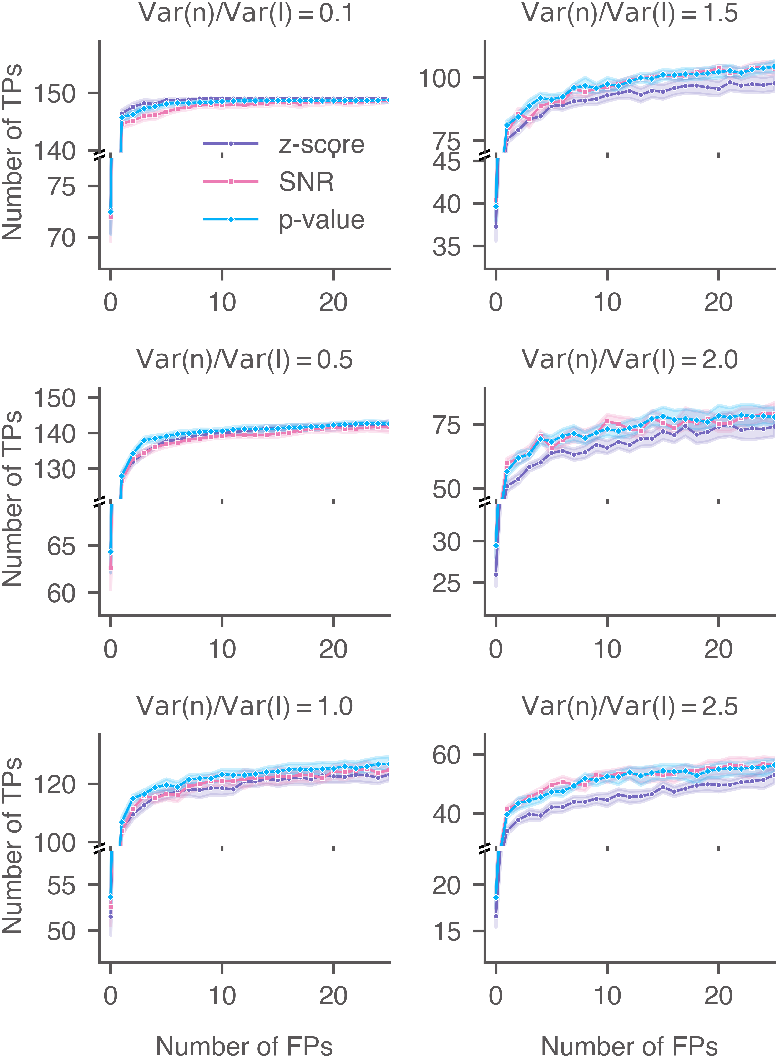
Performance of 2DTM metrics when searching for 60S in yeast lamellae with additional Gaussian noise. ROC curves of the 2DTM SNR, z-score, and p-value for image `150_Mar12_12.28.45_165_0.mrc` with varying levels of Gaussian noise. For each noise level, nine images were generated with random Gaussian noise.

As the variance of the Gaussian noise was further increased (Var(*n*)*/*Var(*I*) *≥* 1.0), the accuracies of the 2DTM z-score experienced a rapid decline in contrast to the other two metrics. The performance of the 2DTM p-value closely mirrored or slightly exceeded that of the SNR at low FPR ranges. In this case, the high levels of Gaussian noise blurred the high-resolution features, forcing detection to primarily rely on the remaining low-resolution signal. The performance of the 2DTM p-value and SNR converged when the noise level was 2.5 (Var(*n*)*/*Var(*I*) = 2.5).

In this test, we imposed an additional constraint during the p-value computation, requiring that both quantile-normalized features exceed 0.5 (*x*_1_ *>* 0.5 and *x*_2_ *>* 0.5 in Eq. 31). This criterion ensures that both SNR and z-score must be high for a true target. An interesting observation is that the features from contamination (dark regions in Fig. 10A), which could potentially lead to false positives when using a *blob detector* solely based on low-resolution signal, were correctly excluded by the 2DTM p-value despite its sensitivity to low-resolution signal.

In summary, introducing Gaussian noise to experimental images from yeast cells, coupled with the curation of control datasets based on the distribution of 2DTM z-scores, enables the evaluation of the 2DTM p-value across varying noise levels. Our findings show that while the z-score performs well for targets such as ribosomes, the p-value is robust under conditions of increased noise.

## 4. Discussion

Detecting biological molecules and complexes in low-contrast cryo-EM images is an important step for determining their molecular structures *in situ* and understanding the mechanisms of biological processes. Previous works have demonstrated that accurate determination of target locations and orientations can be achieved using 2DTM, with high-resolution structures as templates and sampling of poses on a tight grid. 2DTM offers a way to study macromolecular assemblies in the broader context of a cell, taking advantage of the increasing number of high-resolution structures available for templates.

Building on the success of 2DTM in locating and distinguishing larger molecular species in cells, our goal in this paper is to improve 2DTM to detect more challenging targets, especially those that are smaller and aspherical. We show that the outputs of 2DTM, namely the 2DTM SNR and 2DTM z-score, offer complementary information for target detection. By integrating data from both metrics, we introduce a novel 2DTM metric, the 2DTM p-value, which improves the detection of previously unexplored targets, such as clathrin monomers. Our results show that the performance of the 2DTM p-value is robust across diverse imaging conditions. Furthermore, we have established a general framework for combining multiple metrics of varying scales into a new “metafeature”, and developed a probabilistic model for target detection in both purified and cellular cryo-EM images using 2DTM. The approach used to construct the 2DTM p-value is not limited to applications in 2DTM; it can easily be extended to applications in cryo-electron tomography (cryoET), including the development of a detection likelihood model that utilizes multiple metrics derived from 3D template matching (3DTM) (Xue *et al*., 2022; Cruz-Leon *et al*., 2023; Maurer *et al*., 2024).

### Determining a 2DTM p-value threshold for target

In situations where labeled data are unavailable, determining an appropriate threshold for target detection based on the 2DTM p-value is crucial. One approach is to calculate the adjusted p-values for multiple comparisons using the Benjamini-Hochberg procedure (Benjamini & Hochberg, 1995). Subsequently, the quantile of the adjusted p-values corresponding to an estimated number of true positives can be identified. This quantile then serves as the classification threshold. Unlike the 2DTM z-score, which relies on a uniform threshold calculated from the number of search locations, the p-value learns from the signal and noise distribution unique to each 2DTM search. We provide an example of using this method to calculate the 2DTM p-value for a noisy image from the yeast lamella dataset (Appendix Fig. 22), where Var(*n*)*/*Var(*I*) = 0.5. In this example, we identified 166 out of 176 targets.

### Molecular weight and shape jointly affect target detection

Our results in Section 3.1 and 3.2 demonstrate that detecting small and aspherical targets using either 2DTM SNR or z-score alone is particularly challenging due to the significant overlap between true and false targets in both metrics.

The effectiveness of the 2DTM SNR is heavily influenced by the molecular weight of the target and the projected density distribution of the target across the image. Specifically, when the targets are small or have a dispersed projected density, the 2DTM SNR tends to be low, making accurate detection more difficult. Conversely, for targets viewed edge-on with a high projected density, the SNR is higher, but this also increases the risk of misalignment.

The 2DTM z-score works well for detecting spherical targets, where the correlation peak is relatively sharp due to the effective subtraction of rotationally invariant components. This characteristic is a result of the z-score transformation, which was introduced as an “*a posteriori* “ correction of camera characteristics in large imaging datasets (Afanasyev *et al*., 2015) and previously applied in 2DTM to successfully normalize the spurious correlations generated from low-resolution matches (Rickgauer *et al*., 2017; Rickgauer *et al*., 2020). However, it is less reliable for aspherical targets. In these cases, the z-score map tends to have a less clean background and may lead to false positives or false negatives, as in the tubulin patch example. When combining these factors, particularly in the context of small and aspherical particles, the overlap between false and true populations poses a significant challenge for accurately separating and classifying targets. However, more true targets can be recovered by utilizing a “metafeature” like the 2DTM p-value that integrates information from the 2DTM SNR and z-score.

### The 2DTM p-value is robust regardless of image and target characteristics

The 2DTM p-value combines information from the SNR and z-score, providing a more robust metric than either alone. It ensures optimal target detection regardless of the signal characteristics in the image, whether dominated by low- or highresolution features. In many applications of 2DTM (Rickgauer *et al*., 2017; Lucas *et al*., 2021), suppressing low-resolution signal from the background improves overall precision. However, theoretically, the low-resolution signal of the target itself should aid in target detection. The challenge lies in developing a method that accurately leverages the target’s low-resolution signal without losing the ability to distinguish true and false positives when the image contains strong low-resolution contrast.

While the 2DTM p-value incorporates correlations arising from the rotationally invariant or lower resolution components between template and targets, it largely avoids incorrect low-resolution features that may be present in cellular cryo-EM images. Our findings in Section 3.4 show that even in images where incorrect low-resolution features can strongly bias the 2DTM SNR, the 2DTM p-value still out-performs the z-score. This highlights the potential of using the 2DTM p-value for target detection in native cells, even in the presence of cellular background noise, such as membranes and other molecules.

In this work, we present extensions of 2DTM applications specifically targeting aspherical targets, demonstrating the versatility of the 2DTM p-value in diverse experimental scenarios. Using examples of tubulin patches and clathrin monomers, we demonstrate the advantage of using the 2DTM p-value when the 2DTM SNR and z-score may fall short.

### Future improvement

#### Better whitening filter

In 2DTM, we apply a global whitening filter based on the power spectrum of the entire image to whiten the noise spectrum and ensure the accurate matching of signal in the image by the template, both in real and reciprocal space. However, the global whitening filter may not uniformly whiten local areas within the image (Lucas *et al*., 2023). A better whitening strategy that addresses local contrast variations (Roseman, 2003; Roseman, 2004) could enhance the calculation of the z-score, thereby improving the resulting p-value. Another approach to suppress low-resolution structural noises in cellular images is to utilize the phase-only correlation (Horner & Gianino, 1984; Ahmed & Jafri, 2008). However, excluding amplitude information may weaken the overall signal correlation and require multiple passes of template matching.

#### Conformational heterogeneity discrimination

So far, 2DTM metrics have primarily been used for identifying one or a few 3D templates (Lucas *et al*., 2022). However, due to thermal fluctuations, it is expected that an ensemble of conformations will be present within the sample. Additionally, crowded *in situ* environments may result in interactions between the biomolecule and binding partners, potentially causing subtle structural modifications of the template. Consequently, further research is needed to evaluate the discriminatory power of the 2DTM p-value in detecting small structural changes, such as those on the order of a few Å due to thermal fluctuations. Utilizing a library of structures generated through molecular dynamics simulations as templates (Giraldo-Barreto *et al*., 2021; Tang *et al*., 2023*b*; Tang *et al*., 2023*a*) would enable a more comprehensive exploration of the conformational landscape of the molecules present in the image. Deep learning methods that amortize template matching (Dingeldein *et al*., 2024) may be necessary to overcome the challenges of dealing with large structural ensembles, where the computational cost of matching all templates to each image becomes prohibitively high. Additionally, efficient image alignment techniques, such as those using polar coordinates and Fourier-Bessel transformations or SVD-based compression (Rangan *et al*., 2020; Rangan, 2022), can further accelerate the 2DTM angular search.

#### Alternate distribution modeling

Furthermore, the quantile normalization uses a probit-function to transform the marginalized distribution of 2DTM SNRs and z-scores to a standard Gaussian distribution. However, as discussed earlier, the 2DTM z-scores of locations without target signal should follow a GEV distribution with an extended tail compared to a standard Gaussian. False positives might be avoided if we use a GEV distribution to model the z-scores of the false targets instead of the Gaussian distribution. Another caveat in our present approach for computing the 2DTM p-values is that the quantile normalization maintains the ranking of data but does not preserve the distance between data points. Future work could incorporate recent developments in computer vision, such as using quantile-quantile embedding (Ghojogh *et al*., 2021), to allow data transformation while maintaining the local distances among nearby data points.

#### Correlations of correlations

Finally, in our exploration to optimize the use of low-resolution signal, we calculated another correlation value: the correlation between the auto-correlation of a 2D projection and the cross-correlation of the image with that projection. This metric, referred to as the “correlation of correlations” (CoC), was devised to capture additional information beyond the normalized cross-correlation coefficient, particularly in assessing the similarity of the correlation maps (Chen & Grigorieff, 2007). We found that the CoC can be interpreted as the distance between an implicitly chosen latent variable associated with the 3D structure of the template and the image (data not shown). Although a similar p-value combining the CoC and z-score was computed, its performance was found to be lower than the combination of the 2DTM SNR and z-score. We hypothesize that the CoC could be particularly useful in images containing predominantly low-resolution signal, where noise is usually associated with high-resolution signal. Our previous study showed that the product of the CoC and the cross-correlation can serve as a better particle picker in single-particle datasets (Chen & Grigorieff, 2007). However, additional research is required to determine the appropriate weighting and integration of the CoC for template matching in more crowded environments.

## Code availability

In *cis*TEM, the implementation of the method described in this paper is provided by the program `calculate template pvalue`.

## Appendix A

### Differences between 2DTM SNR and z-score

In the main text, we have shown that *r*(*i, j, τ*) is an affine transformation of the logarithm of the probability *P* (*Y* |*i, j, τ*) of observing the image *Y*, given a single particle at location *i, j* and orientation *τ*,

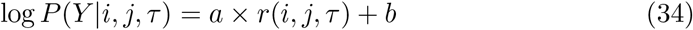

where *a* = *σ*_*Y*_ *σ*_*t*_′ and 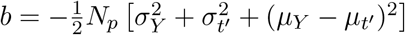.

Similarly, the SNR, *r*(*i, j*), is up to the same affine transformation, the maximum log-likeihood log *P* (*Y* |*i, j, τ*), optimized over *τ*. For these reasons, we expect the SNR to highlight locations and orientations that correspond to peaks in the posterior likelihood of observing the data, given the position *i, j* and orientation *τ*.

By contrast, the z-score is designed to highlight only those locations where the maximum-likelihood orientation is significantly more likely than other orientations for that location. In practice, the z-score *z*(*i, j*) often approximates a measurement of the ‘Fisher information’ at that location – i.e., a measurement of how tightly the likelihood is peaked around the maximum *r*(*i, j, τ*) as a function of orientation.

To understand why this might be the case, let us assume that *r*(*i, j, τ*) adopts a roughly Gaussian profile around its maximum value

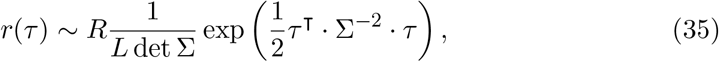

with *L* = (2*π*)^(3*/*2)^. Then we see that

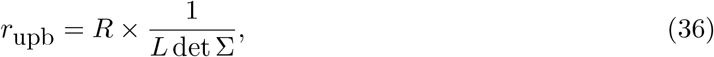

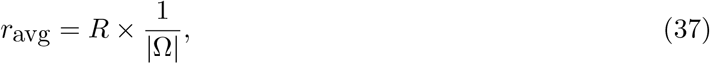

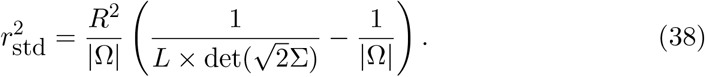

According to the definition of the z-score, and assuming |Ω| is relatively large compared to det Σ, we have

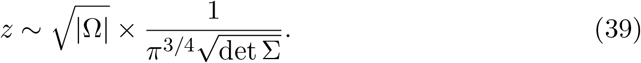

The Fisher information regarding perturbations in *τ* is:

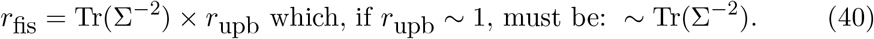

If *r* is roughly isotropic, the eigenvalues of Σ will be close to one another, and the Fisher information will once again be close to a (fixed) power of the z-score.

In summary, we expect that peaks in the SNR map should roughly correspond to locations (and orientations) that are local peaks in the log-likelihood of observing the data, given that a true particle is being imaged. Note that the SNR does not require any particular orientation to be more likely than any other and will pick out locations that have broad peaks with respect to orientation. Conversely, we expect that peaks in the z-score should roughly correspond to locations (and orientations) that have high amounts of Fisher information regarding orientation – that is, locations where a particle can be unambiguously aligned to the image. In practice, these two measurements are not entirely redundant and can even complement one another; as we have shown in the Result section, the SNR and the z-score can be combined into an even more informative metric.

In Appendix Fig. 12, we calculated the normalized cross-correlations between the simulated particles shown in Fig. 3 and a series of 2D projections, where the angles *ψ* and *ϕ* were kept the same as those of the simulated particles, while the polar angle *θ* was uniformly sampled between 0^°^ and 180^°^. The polar angles from the simulated particles, *θ*_0_, were labeled. As shown in the plots, *r*(*i, j, θ*) exhibits a sharp, unimodal peak near *θ*_0_ for all three particles.

## Appendix B

### Zernike decomposition of cryo-EM density maps

The Zernike decomposition of a 3D density map *V* (**r**) can be expressed as:

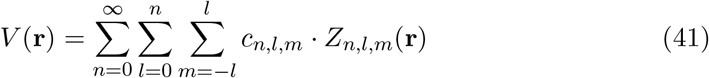

where:

- **r** = (*x, y, z*) is the position vector in 3D space.
- *c*_*n,l,m*_ are the Zernike coefficients, representing the contribution of each Zernike polynomial to the density map.
- *Z*_*n,l,m*_(**r**) are the Zernike polynomials, defined as the product of radial and angular components:

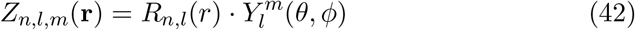
- *R*_*n,l*_(*r*) is the radial polynomial, and 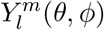is the spherical harmonic function.

Here, *r, θ*, and *ϕ* are the spherical coordinates related to the Cartesian coordinates (*x, y, z*).

The rotationally invariant components are those that are associated with the coefficients where *m* = 0, and the rotationally variant components involves terms where *m≠* 0:

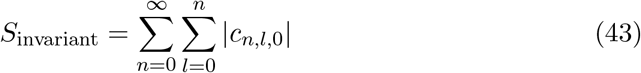

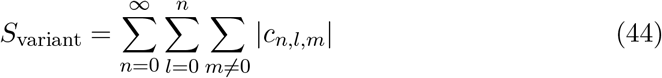

To quantify the asphericity of apoferritin and the modified microtubule, we bin the density map to 64 *×* 64 *×* 64 and calculate the Zernike decomposition with order 120 using codes from the GitHub repository `zernike3d` (Bayly-Jones, 2024).

## Appendix C

### Modeling the 2DTM z-scores using the generalized extreme value distribution

### C1. Introduction of the generalized extreme value distribution

The family of generalized extreme value (GEV) distributions is frequently used to model the maxima (or minima) of a large set of random variables. The GEV distribution combines the Gumbel, Fréchet, and Weibull distributions into a single family with a common cumulative density function (CDF) given by:

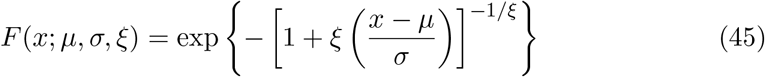

where:

- *µ* is the location parameter,
- *σ >* 0 is the scale parameter,
- *ξ* is the shape parameter.

Note that 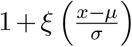 should be greater than zero. The probability density function (PDF) is:

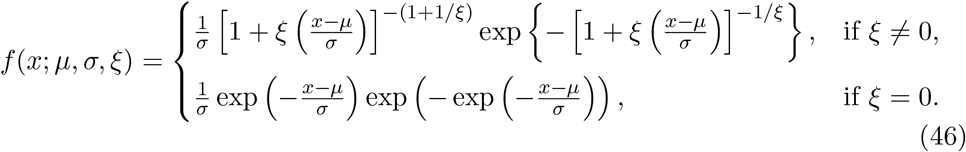

When *ξ* is zero, the distribution is simplified to the Gumbel distribution.

#### C2. Modeling the z-score map using the GEV distribution

The cross-correlations *r*(*i, j*) at different locations in the cryo-EM image can have different statistical behaviors: (a) for locations without targets and dominated by shot noise, the correlations are primarily affected by noise, leading to overall low values and possibly higher variations; (b) for locations with background noise (membranes or dense structures in the cell), the correlations tend to have higher values but still do not depend on orientations; (c) for locations with searched targets, correlations depend on the orientation and can be significantly higher at the correct orientation.

The 2DTM z-score is essentially a scaled maximal value drawn from a large number of correlations. For locations without targets (cases (a) and (b)), the normalization during the z-score calculation removes local variations caused by heterogeneous densities (e.g., membranes and other cellular structures) and varying imaging conditions, aligning the z-scores more closely with the GEV distribution.

We thank the Grigorieff Lab members for the fruitful discussion on this project.

The Flatiron Institute is a division of the Simons Foundation.

---

**Fig. 12.**
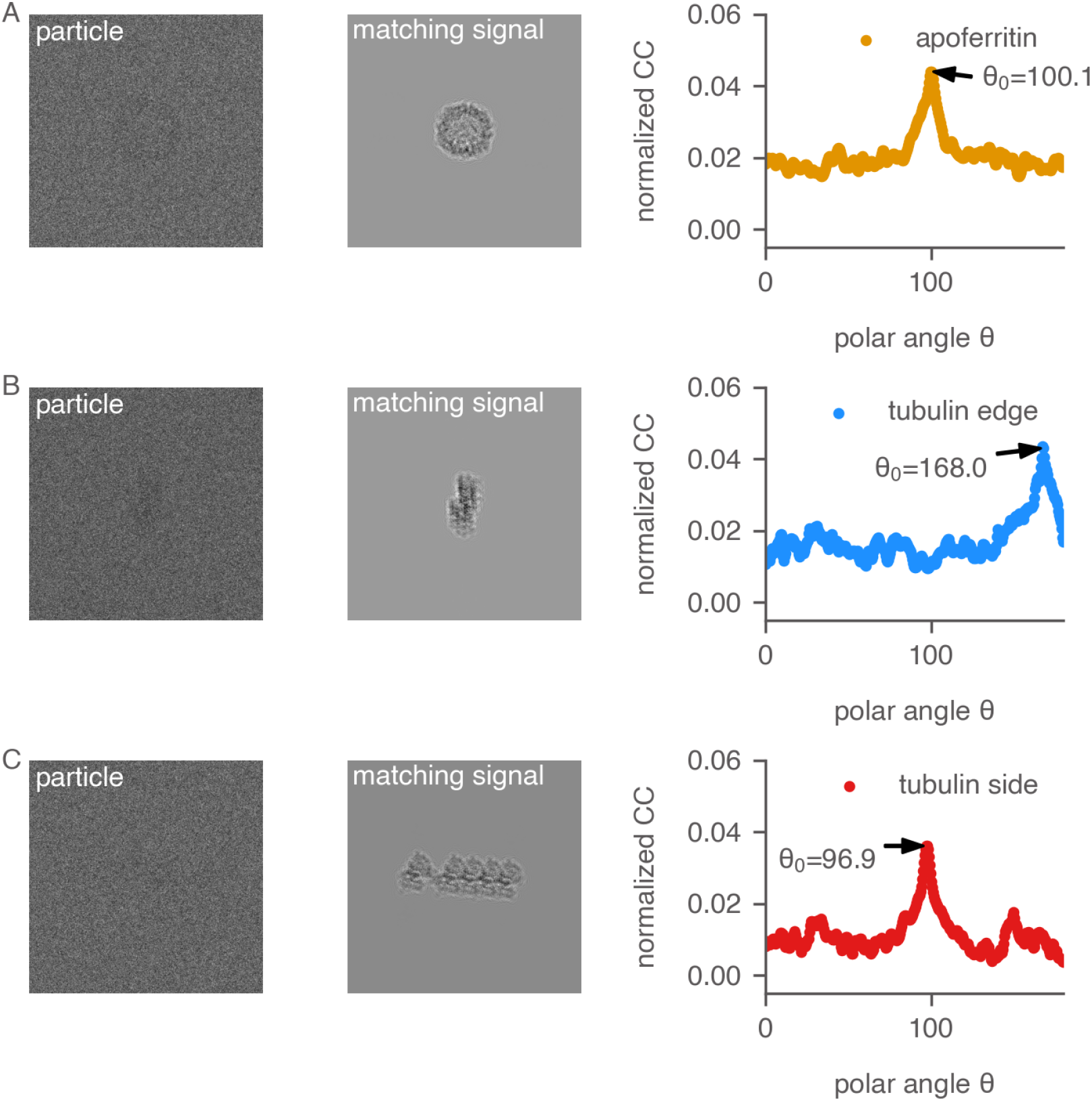
Normalized cross-correlation *r*(*i, j, θ*) between simulated particle images and 2D projections generated at varying *θ*. The 2D projections were generated by keeping the angles *ψ* and *ϕ* equal to those of the simulated particles while uniformly sampling the polar angle *θ* at 0.5^°^ intervals between 0 and *π*. In all three cases, *r*(*i, j, θ*) is sharp around the optimal *θ* = *θ*_0_.

**Fig. 13.**
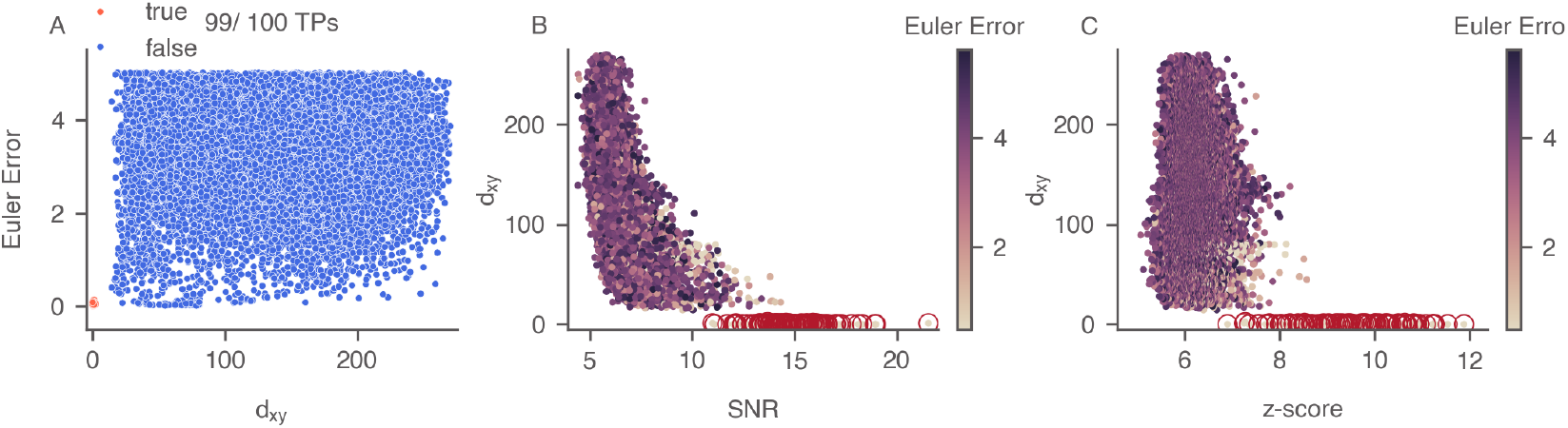
Error distribution for the example tubulin patch montage in Fig. 4A. The angular error was calculated based on the average l2 distances between corresponding points in the two (unit vector) templates after angular transformation. The translational error (*d*_*xy*_) was defined as the distance between the target and the grid center of the closest simulated particle. The cutoffs used for labeling were 7 pixels for *d*_*xy*_ and 0.4 for angular error. (A) Angular error distribution between the 2DTM z-score-derived targets and the ground truth. A total of 26783 local maxima were identified in the z-score map using a local radius of 10 pixels and a threshold of 0. In this example, 99 out of 100 simulated particles were recovered as local maxima. (B) and (C) Distribution of *d*_*xy*_ of the 2DTM-derived targets. True positives are indicated by circles.

**Fig. 14.**
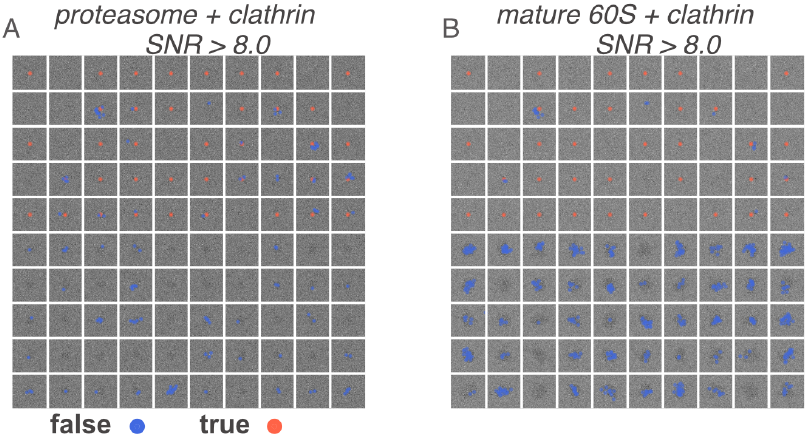
High 2DTM SNR values identify locations that overlap with protein particles. Shown are locations in two example mixed particle montages where the 2DTM SNR is greater than 8.0: (A) a clathrin and proteasome mixture, and (B) a clathrin and mature 60S mixture. This demonstrates that the 2DTM SNR can function as a *blob detector*.

**Fig. 15.**
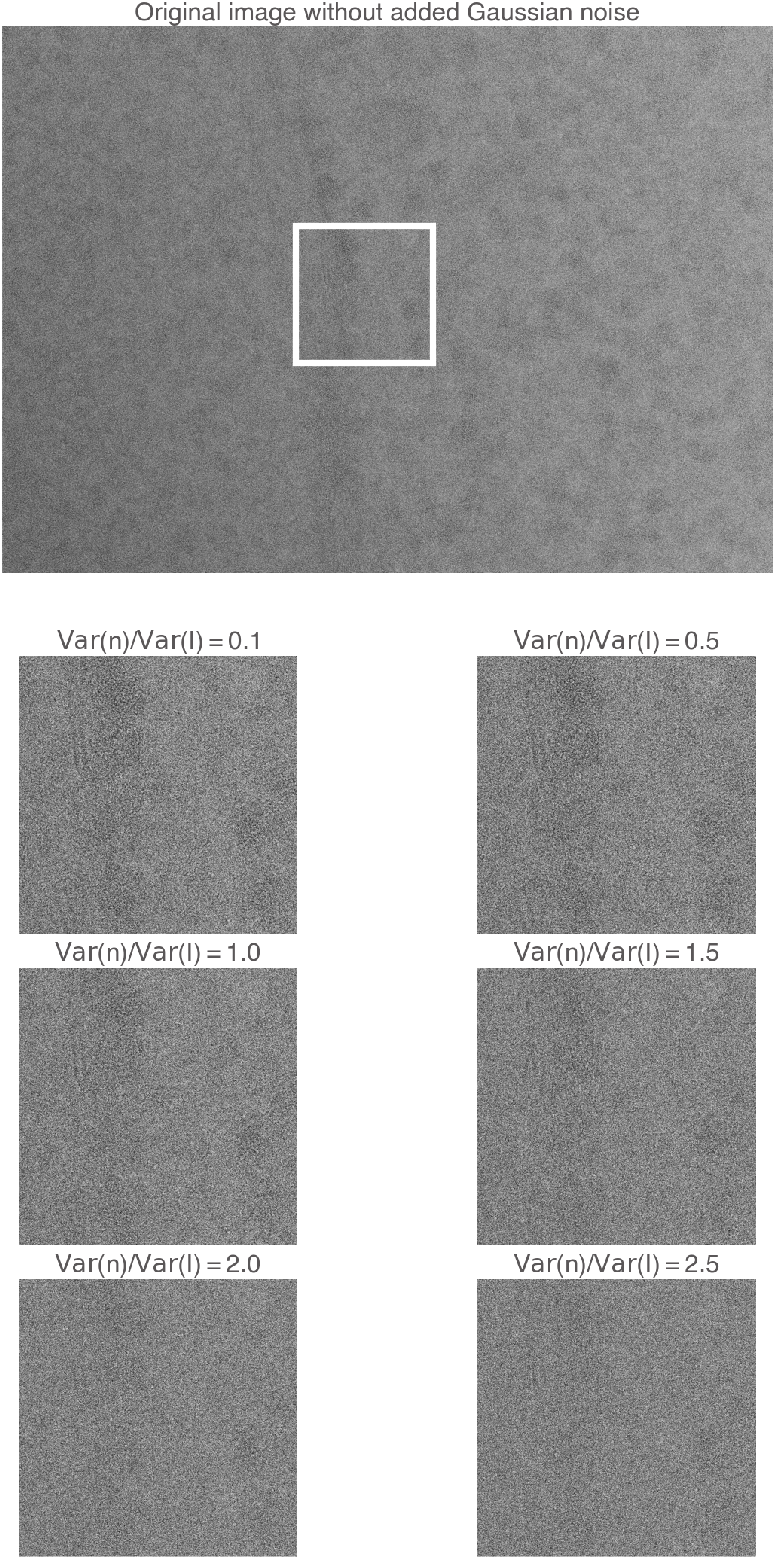
Original yeast lamella image (148_Mar12_12.23.52_161_0.mrc) and images with varying levels of added Gaussian noise. The ratio of added noise to the original image noise ranges from 0.1 to 2.5. The image segment within the white box is shown below.

**Fig. 16.**
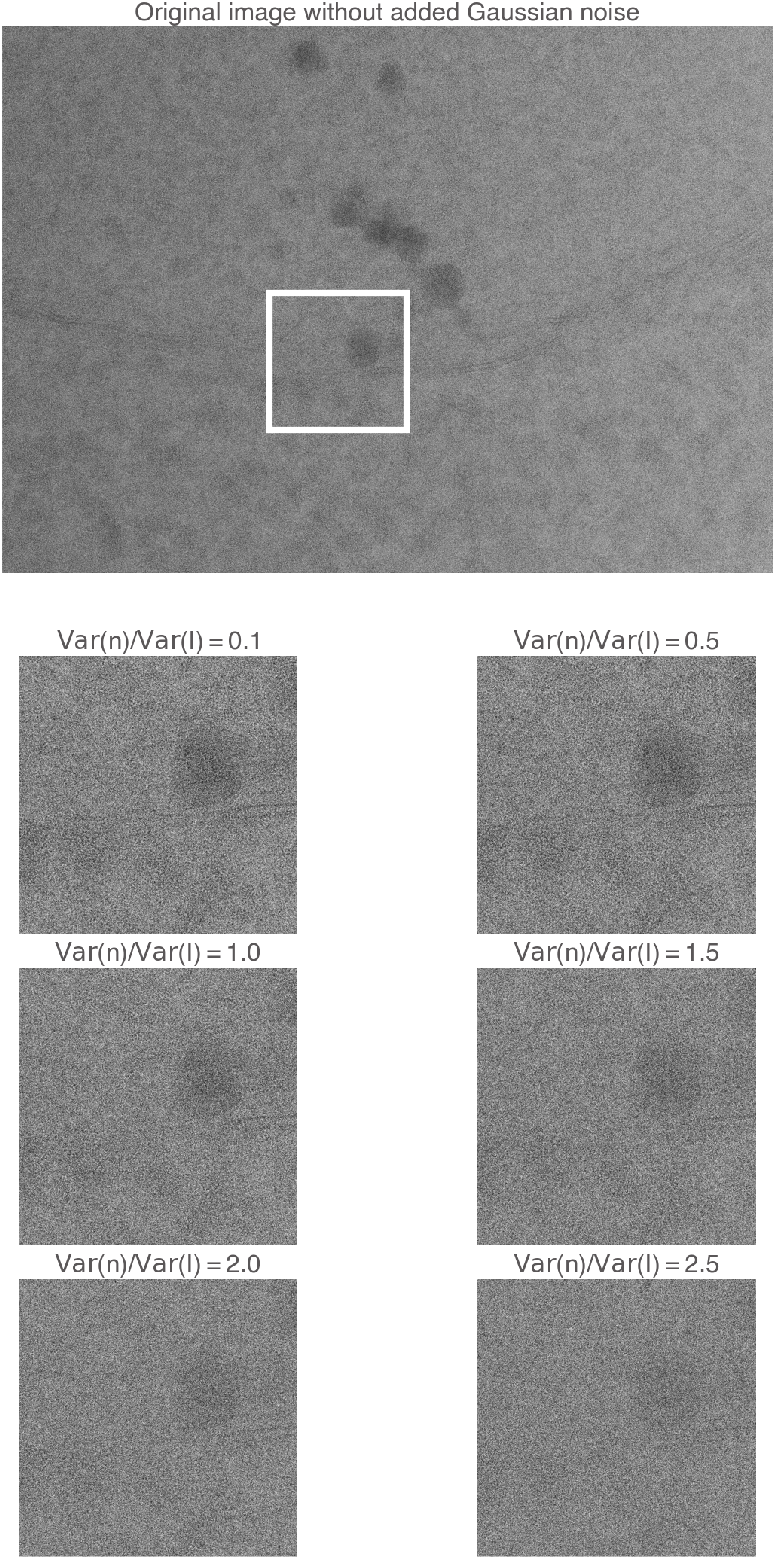
Original yeast lamella image (150_Mar12_12.28.45_165_0.mrc) and images with varying levels of added Gaussian noise. The ratio of added noise to the original image noise ranges from 0.1 to 2.5. The image segment within the white box is shown below.

**Fig. 17.**
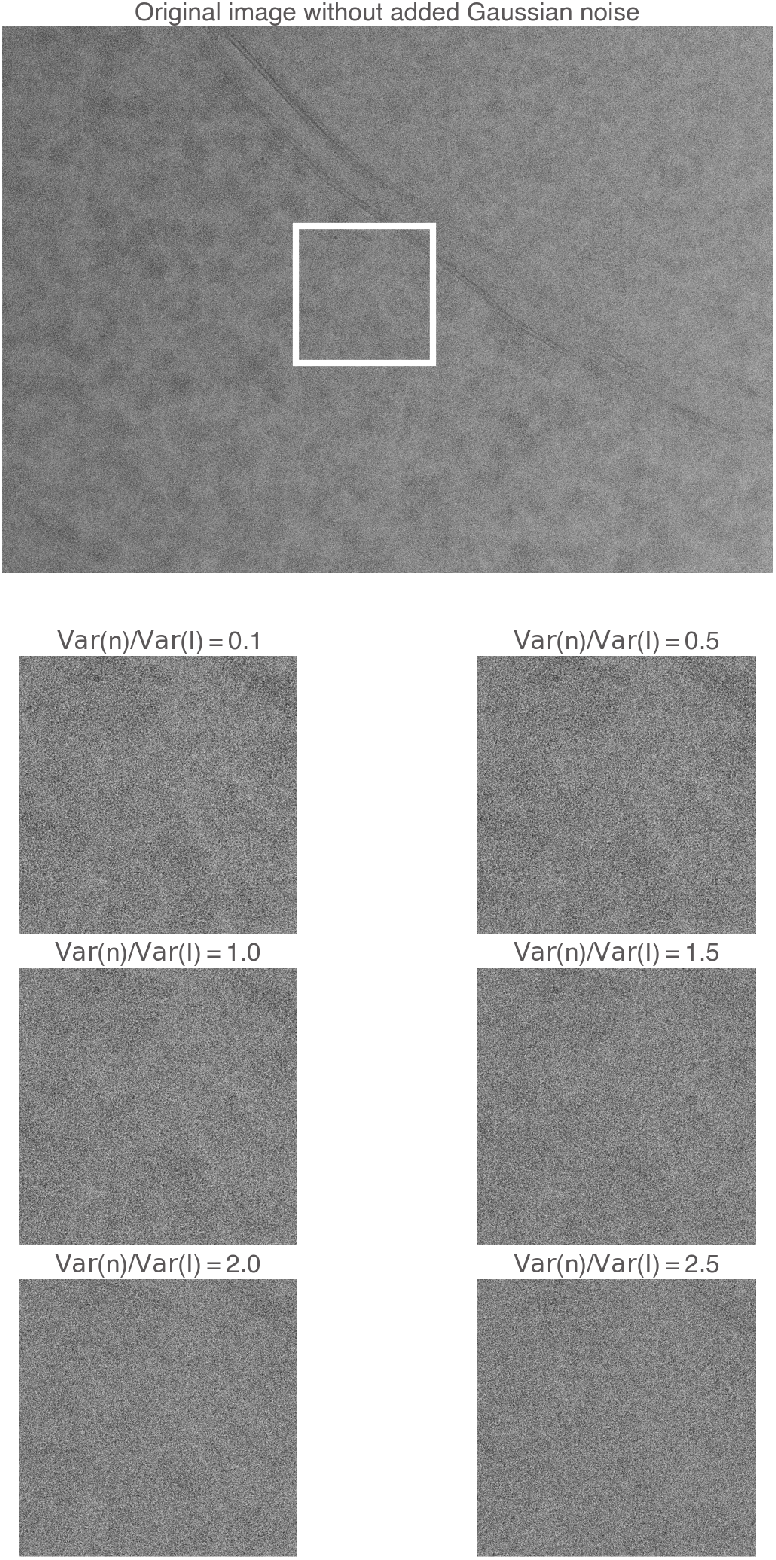
Original yeast lamella image (151_Mar12_12.31.16_167_0.mrc) and images with varying levels of added Gaussian noise. The ratio of added noise to the original image noise ranges from 0.1 to 2.5. The image segment within the white box is shown below.

**Fig. 18.**
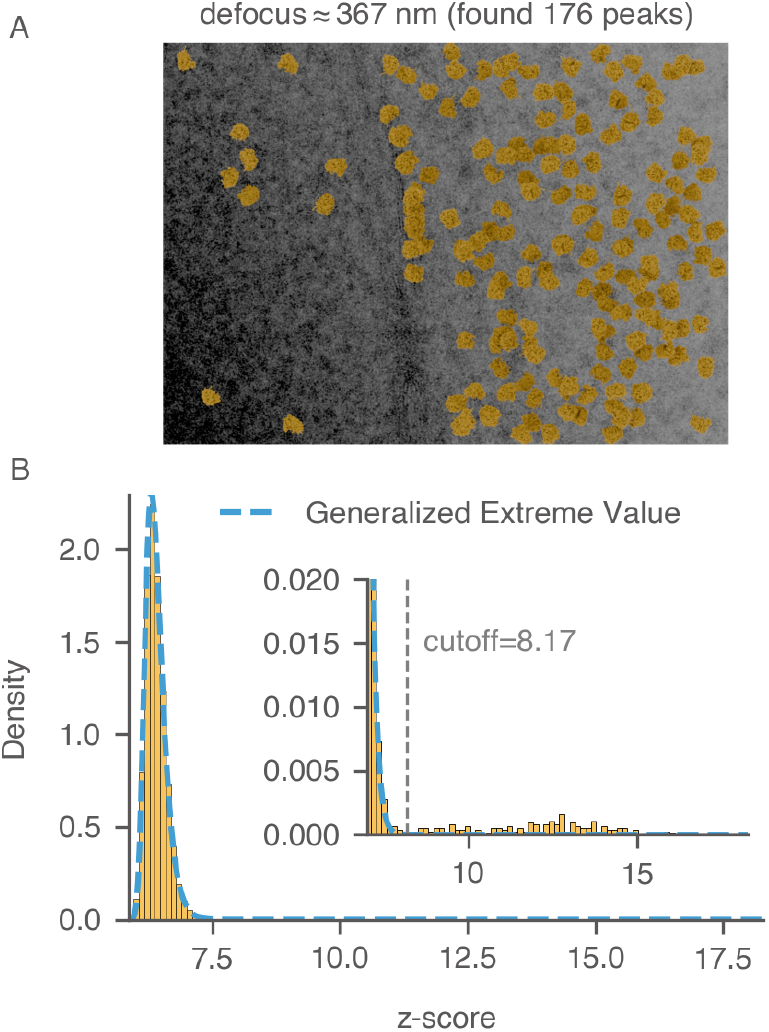
Labeling 60S targets in yeast lamella. (A) Image `148_Mar12_12.23.52_161_0.mrc` from previous work (thickness was estimated to be 73 nm) (Lucas *et al*., 2022). Control peaks are selected based on the threshold determined in (B). Particles are plotted with their 2DTM-derived alignment parameters. (B) Distribution of z-scores across all locations in the image using mature 60S as the template. The dashed blue curve represents the fitted generalized extreme value distribution. The threshold that best separates false matches (bulk) from true matches (tail) is labeled.

**Fig. 19.**
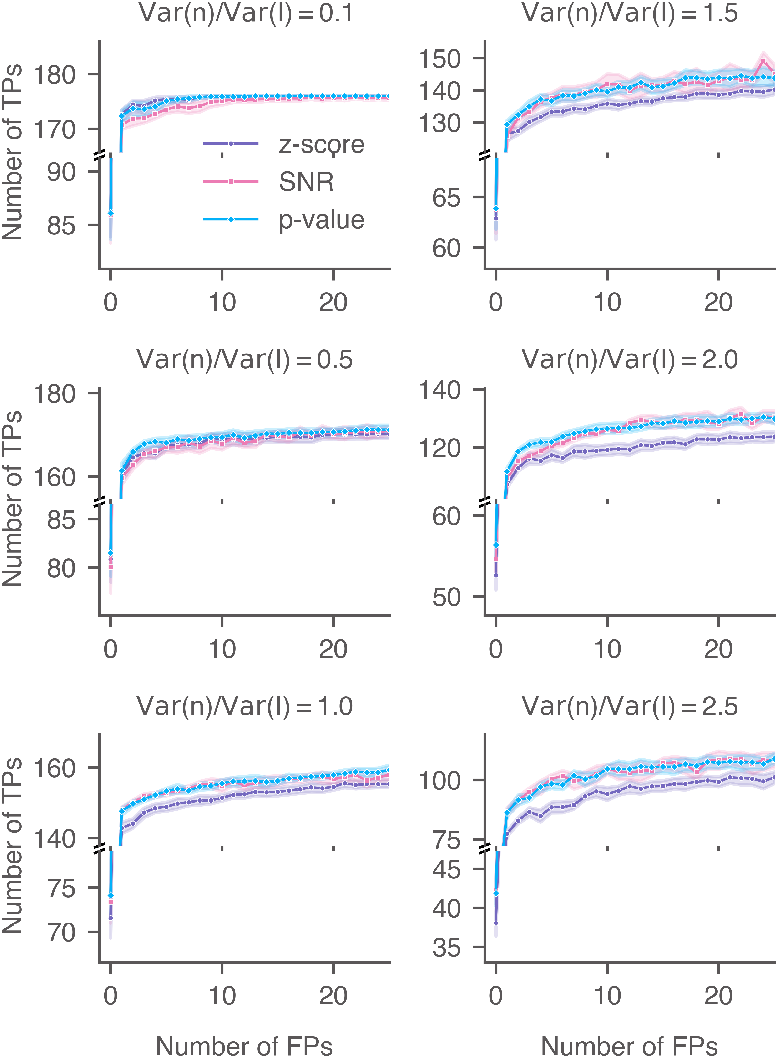
Performance of 2DTM metrics when searching for 60S in yeast lamellae with additional Gaussian noise. ROC curves of the 2DTM SNR, z-score, and p-value for image `148_Mar12_12.23.52_161_0.mrc` with varying levels of Gaussian noise. For each noise level, nine images were generated with random Gaussian noise.

**Fig. 20.**
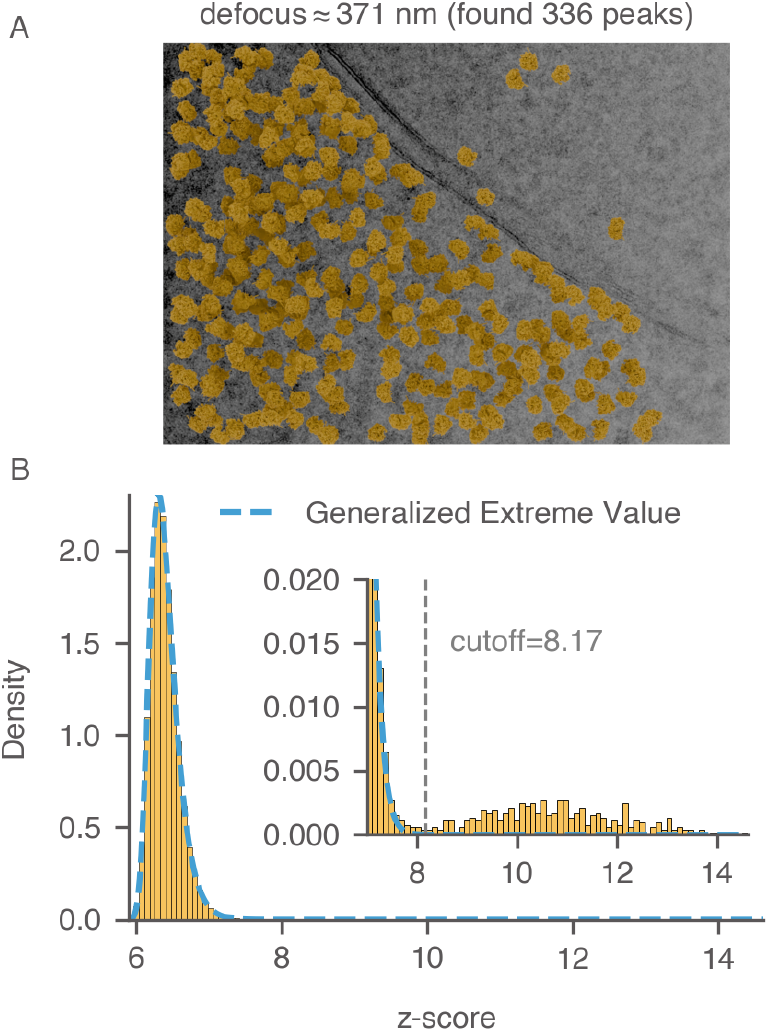
Labeling 60S targets in yeast lamella. (A) Image `151_Mar12_12.31.16_167_0.mrc` from previous work (thickness was estimated to be 114 nm) (Lucas *et al*., 2022). Control peaks are selected based on the threshold determined in (B). Particles are plotted with their 2DTM-derived alignment parameters. (B) Distribution of z-scores across all locations in the image using mature 60S as the template. The dashed blue curve represents the fitted generalized extreme value distribution. The threshold that best separates false matches (bulk) from true matches (tail) is labeled.

**Fig. 21.**
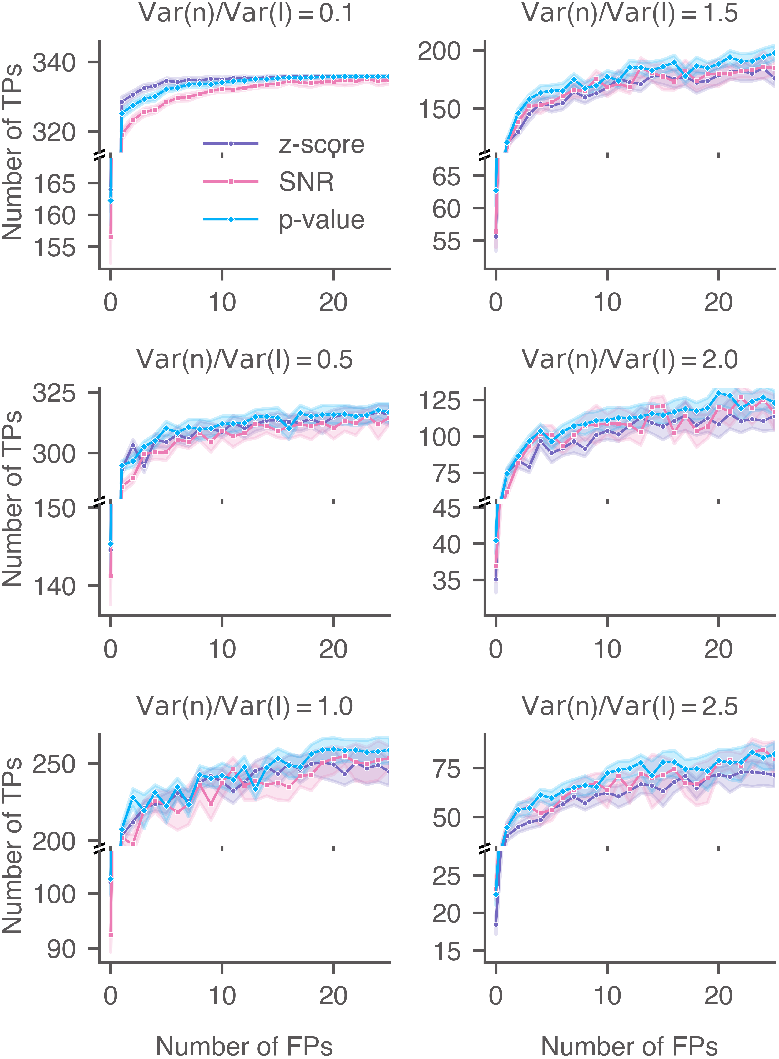
Performance of 2DTM metrics when searching for 60S in yeast lamellae with additional Gaussian noise. ROC curves of the 2DTM SNR, z-score, and p-value for image `151_Mar12_12.31.16_167_0.mrc` with varying levels of Gaussian noise. For each noise level, nine images were generated with random Gaussian noise.

**Fig. 22.**
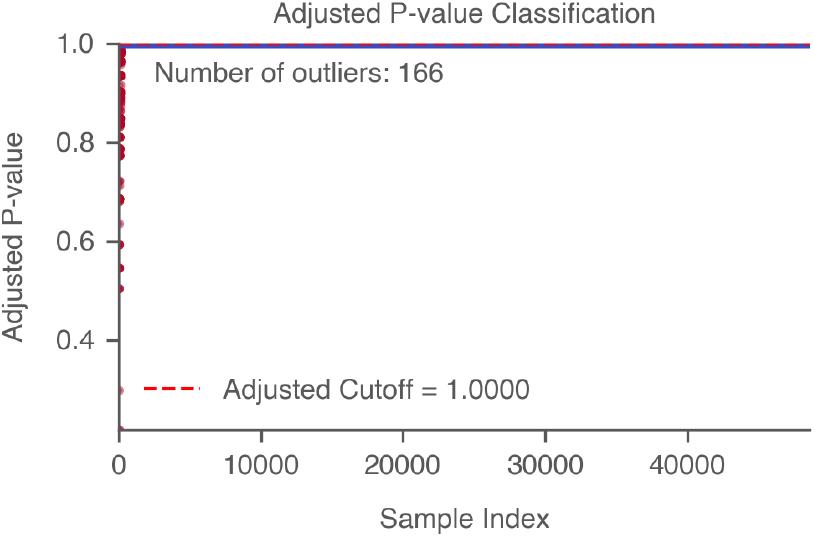
Calculating the 2DTM p-value threshold using multiple hypothesis testing. In this example, we calculated the 2DTM p-values for a noisy image from the yeast lamella dataset (`148_Mar12_12.23.52_161_0.mrc`) with a noise variance ratio of 0.5. We estimated 200 true positives and calculated the adjusted p-values using the Benjamini-Hochberg procedure to control the false discovery rate (FDR) at an alpha value of 0.05. Using this method, we identified 166 out of 176 targets.

**Table 1.** Quantification of target’s asphericity.

| Target | order | $S_{\text{invariant}}$ | $S_{\text{variant}}$ | $\frac{S_{\text{invariant}}}{S_{\text{invariant}} + S_{\text{variant}}}$ |
| --- | --- | --- | --- | --- |
| apoferritin | 120 | 0.0020 | 0.0122 | 0.1388 |
| tubulin patch | 120 | 0.0007 | 0.0095 | 0.0722 |
| clathrin monomer | 120 | 0.0007 | 0.0087 | 0.0782 |

**Table 2.** Generalized extreme value parameter fitting of 2DTM z-scores of yeast lamella images.

| image ID | shape ($c$) | location ($\mu$) | scale ($\sigma$) |
| --- | --- | --- | --- |
| 148 | 0.024 | 6.311 | 0.16 |
| 150 | 0.021 | 6.309 | 0.159 |
| 151 | 0.026 | 6.309 | 0.16 |

